# Minimal DNA Electron Transfer Catalysts Switched by a Chaotropic Ion

**DOI:** 10.1101/784561

**Authors:** Tanner G. Hoog, Matthew R. Pawlak, Lauren M. Aufdembrink, Benjamin R. Bachan, Matthew B. Galles, Nicholas B. Bense, Katarzyna P. Adamala, Aaron E. Engelhart

## Abstract

Here we demonstrate that a DNA nanodevice can perform switchable electron transfer. The nanodevice is comprised of two strands, one of which can be selectively switched between a G-quadruplex and duplex or single-stranded conformations. In the G-quadruplex state, it binds the cofactor hemin, enabling peroxidase activity. This switching ability arises from our discovery that perchlorate, a chaotropic Hofmeister ion, selectively destabilizes duplex over G-quadruplex DNA. By varying perchlorate concentration, we show that the device can be switched between states that do and do not catalyze electron transfer catalysis. State switching can be achieved in three ways: thermally, by dilution, or by concentration. In each case, when operated in the presence of the cofactor hemin, the device catalyzes electron transfer in only the G-quadruplex state.

## Background

Biomolecules are highly attractive as potential programmable electron transfer catalysts and bioelectronics. A range of techniques have been employed, including microbial systems, protein-directed assembly of metal clusters, metallated base pairs, biopolymer-directed assembly of nanoparticles, and minimal peptides.^1–6^ With 0.34 nm spacing between nucleobases and exquisite, atomic-precision self-assembly directed by nucleobase hydrogen bonding, nucleic acids are an especially attractive means by which to develop programmable electron transfer catalysts. DNA possesses inherent conductivity; a hole generated by a single-electron oxidation can propagate through a double helix. However, the utility of DNA as a general-purpose conductor is limited by the fact that each nucleobase possesses a different oxidation potential.^7^ As a result, a hole generated by single electron oxidation of DNA will migrate to a guanine, the most readily oxidized nucleobase, generating guanine radical cation, which in turn can undergo a range of chemical transformations, generating damage lesions such as 8-oxo-7,8-dihydroguanine and 2,5-diamino-4H-imidazol-4-one.^8^ One means by which DNA can perform electron transfer and avoid such damage, enabling multiple turnover of electron transfer, is by employing a prosthetic group, as many enzymes that perform electron transfer in nature do. Hemin, one such prosthetic group, binds selectively to a noncanonical structure formed by G-rich sequences of DNA, termed a G-quadruplex, which activates it to perform electron transfer.^9–11^ By binding hemin, the same prosthetic group employed by natural redox enzymes, such as peroxidases and Cytochrome P450s, DNA can perform electron transfer in a biomimetic fashion while mitigating the oxidative damage issues associated with attempting to use it as a classical conductor.^12^

Sequences of DNA and RNA that can exhibit either G-quadruplex or non-G-quadruplex (e.g., unpaired or Watson-Crick base paired) structures depending on context are a ubiquitous feature of life. These sequences are actively remodeled in living organisms.^13–15^A bioinspired chemical system based on these phenomena that enabled reversible, programmable structure switching would afford a powerful tool for dynamic electron transfer behavior in DNA nanostructures. We thus sought to develop a chemical system to switch the same DNA between secondary structure states, reasoning that this would combine the speed and repeated reversibility of pH-switchable DNA nanomotors with the compatibility observed in static pH strand-exchange based systems.^16–21^

Base pairs have ca. one half the solvent-buried hydrophobic surface area of G-quartets, and ions exert Hofmeister effects by interactions with the surface of biopolymers.^22^ Similarly, G-quartets coordinate dehydrated ions, and high-salt solutions influence biopolymer folding by osmotic effects.^22,23^ We thus reasoned that a DNA duplex and G-quadruplex would exhibit differential destabilization by chaotropes. Here, we demonstrate that perchlorate, a chaotropic anion that is ubiquitous in lithium-ion battery electrolytes, is a selective denaturant for duplex vs. quadruplex DNA. We have exploited this phenomenon to develop a minimal electron transfer catalyst made of DNA that can be switched between three states: a duplex, a G-quadruplex, and a single-stranded state. We show that this switching can be performed thermally, by dilution, or by concentration. We show that the device can be switched over 100 times without degradation, and that it can perform multiple-turnover electron transfer catalysis by binding hemin and catalyzing electron transfer in the G-quadruplex state (**Figure 1**).

**Figure 1.**
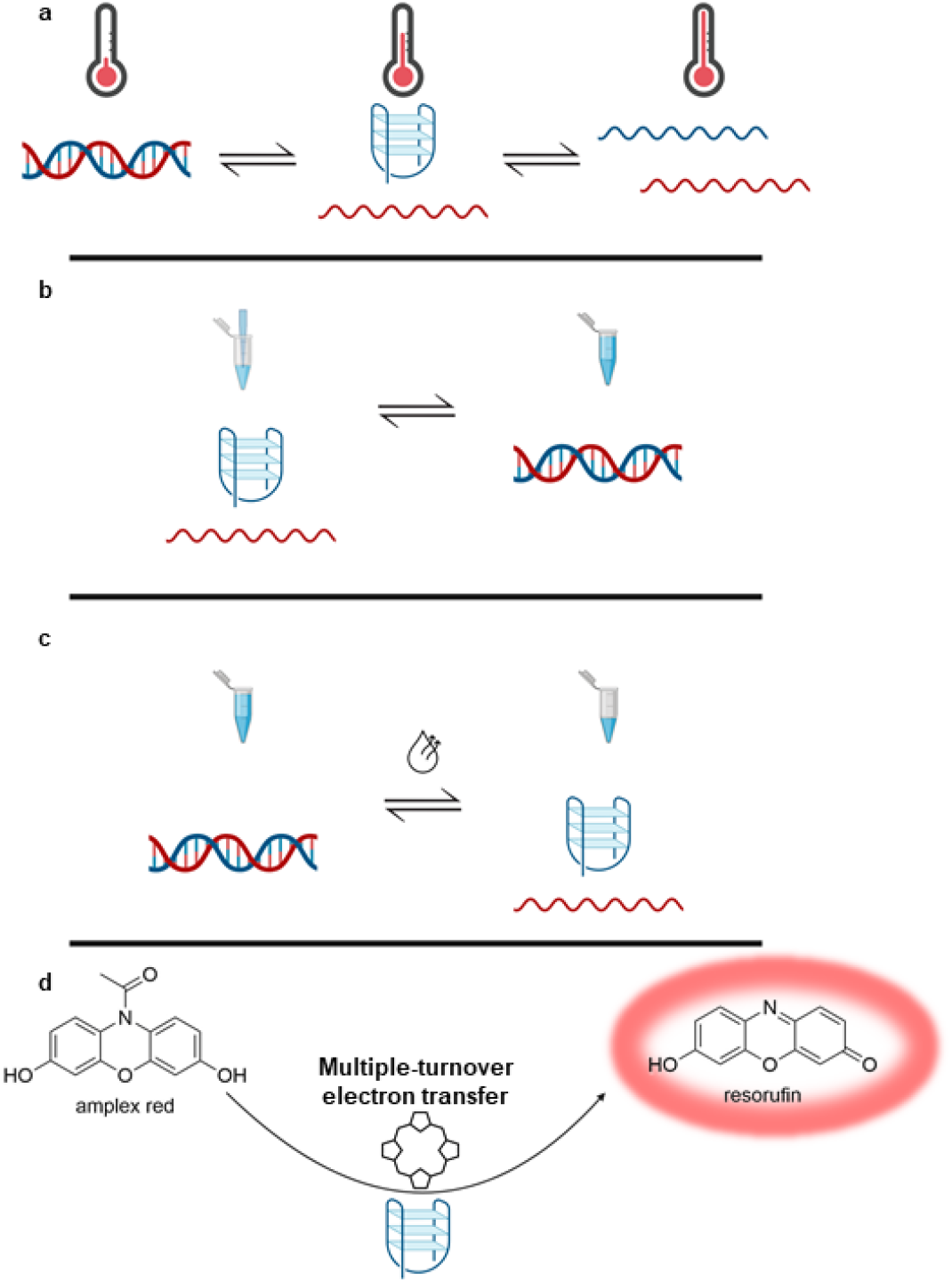
Mechanisms of switching the catalyst: thermally (panel **a**), by dilution (panel **b**), and by concentration (panel **c**). In intermediate perchlorate concentrations, the catalyst can be switched thermally (**a**) through three states. At low temperatures (left), it exists as a duplex; at intermediate temperatures, it exists as a G-quadruplex (center); and at high temperatures, it is single-stranded (right). At high perchlorate concentrations, it exists in only two states - only the G-quadruplex (center) and single-stranded (right) states. The catalyst can also be switched between two states by varying concentration. In high-perchlorate solution, the device exists as a G-quadruplex, while in low-perchlorate solution, it forms a duplex. Thus, by diluting a high-salt solution (panel **b**), the device can be switched from a G-quadruplex to a duplex; and by removal of water from a low-salt solution (panel **c**), the device can be switched from a duplex to a G-quadruplex. In each case, the G-quadruplex form of the catalyst is competent to perform signal amplification by catalyzing multiple-turnover electron transfer in the presence of hemin (panel **d** and **Figure 4**).

## Results

Förster Resonance Energy Transfer (FRET) is a means of nonradiative energy transfer mediated by dipole-dipole coupling of two chromophores.^24^ The r^6^ distance dependence of this phenomenon affords exceptionally sensitive response to the structural state of the system in use.^24^ We designed two FRET reporter systems, **G4-Dark** and **Duplex-Dark** (**Figure 2a** and **Table 1**), to allow readout of their folding state. **G4-Dark** was comprised of equimolar amounts of **Fluorescein-G4-Quencher,** a DNA sequence that could form a G-quadruplex, and **G4Comp**, its Watson-Crick complement. **Fluorescein-G4-Quencher** was 5’-labeled with a fluorescein tag and 3’-labeled with a quencher. Thus, this system would exhibit fluorescence when **Fluorescein-G4-Quencher** was either unfolded or hybridized to **G4Comp.** When **Fluorescein-G4-Quencher** was folded into a G-quadruplex, fluorophore and quencher would be brought into spatial proximity and quenched.

**Figure 2.**
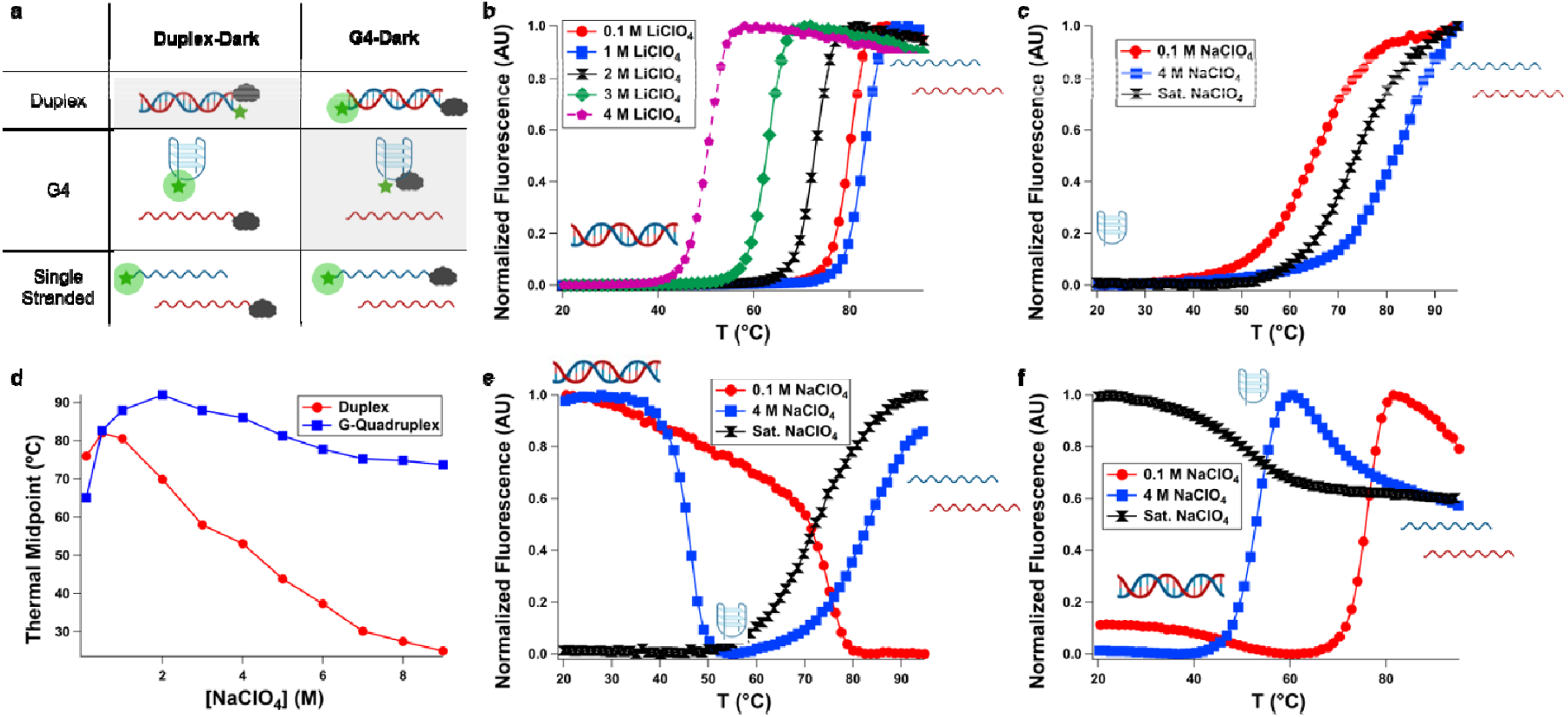
Systems employed in fluorescence-monitored thermal denaturation experiments (panel **a**). **Duplex-Dark** (left column of panel **a**) consists of a 5’-fluorescein labeled strand (green star) and a 3’-Iowa Black FQ (grey cloud) strand and can exist in three states: A quenched duplex, a dequenched G-quadruplex and single strand, and a dequenched set of two single strands. **G4-Dark** (right column of panel **a**) consists of a dual-labeled (5’-fluorescein, 3’-Iowa Black FQ) strand and its complement and can also exist in three states: A dequenched duplex, a quenched G-quadruplex and single strand, and a dequenched set of two single strands. When **Duplex-Dark** is operated in LiClO_4_ solution, only duplex and single-stranded states are accessible. This transition is destabilized with increasing perchlorate (panel **b**). When only the **Fluorescein-G4-Quencher** component of **G4-Dark** is operated in NaClO_4_ solution (panel **c)**, the G-quadruplex-forming strand is significantly less destabilized by perchlorate (panel **d**), in contrast to the duplex, which exhibits ca. linear destabilization with increasing NaClO_4_ concentration above 1 M (panel **d**). As a result of this differential stability, **G4-Dark** (panel **e**) and **Duplex-Dark** (panel **f**) can be switched thermally between duplex, G-quadruplex, and single-stranded states, and the temperatures at which these transitions occur can be tuned by varying the concentration of NaClO_4_. In low perchlorate (0.1 M), only the duplex-to-single stranded transition is observed (red circles). In intermediate perchlorate (4 M), the device transitions between duplex (at low temperature), G-quadruplex (at intermediate temperature), and single-stranded (at high temperature) states (blue squares). In high perchlorate (saturated, ca. 9 M), the device exists only in the G-quadruplex and single-stranded states (black hourglasses).

**Table 1.**
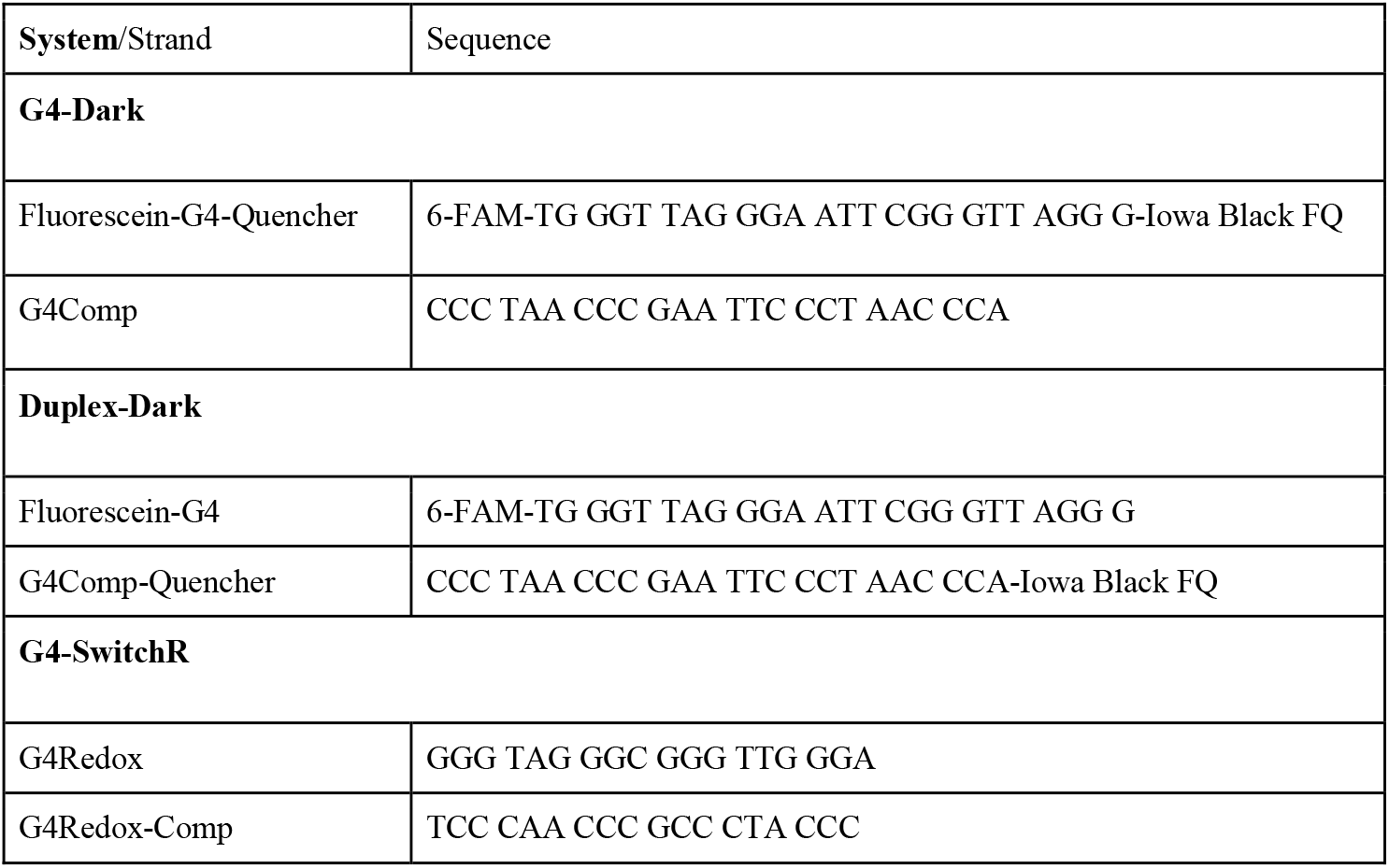
Sequences employed in this study.

**Duplex-Dark** was comprised of equimolar amounts of strands of the same sequences as we employed in **G4-Dark** with rearranged reporters. **Fluorescein-G4,** a 5’-fluorophore-labeled sequence that could form a G-quadruplex, and **G4Comp-Quencher, a** 3’ quencher-labeled sequence. In this system, the fluorescein reporter exhibits fluorescence when **Fluorescein-G4** is either folded into a G-quadruplex or is unfolded. Upon hybridization of the two strands to form a double helix, this system would quench.

We first examined the thermal response of the **Duplex-Dark** system in LiClO_4_ solution (conditions in which only duplex and unfolded states are possible, due to the stability trend in G-quadruplexes (K^+^>Na^+^>>>Li^+^) **(Table 2**, **Figure 2b**, **Supplementary Figures 1-6**).^23^ At 100 mM LiClO_4_, **Duplex-Dark** exhibited fluorescence response consistent with a double-stranded to unfolded structural transition at 79.8 °C (**Duplex-Dark** possesses spatially adjacent reporters in the duplex state, resulting in a systematic slight elevation of T_M_ relative to other sequence-related systems with differing tags). As the salt concentration was increased, the thermal midpoint initially increased to a maximum of 84.7 °C in 0.5 M LiClO_4_, above which it decreased to a minimum of 50.1 °C in 4 M LiClO_4_. This is consistent with electrostatic stabilization effects being predominant at lower salt concentrations and Hofmeister ion effects at higher concentration, as has been previously observed.^25^

**Table 2.** Thermal midpoints (°C) of **G4-Dark, Duplex-Dark,** and **Fluorescein-G4-Quencher** in NaClO_4_ solution, and of **G4-Dark** and **Duplex-Dark** in LiClO_4_ solution.

| Strands/[ClO<sub>4</sub>] (M) |  | 0.1 | 0.5 | 1 | 2 | 3 | 4 | 5 | 6 | 7 | 8 | Sat. |
| --- | --- | --- | --- | --- | --- | --- | --- | --- | --- | --- | --- | --- |
| **G4-Dark (Fluorescein-G4-Quencher and G4Comp)/NaClO<sub>4</sub>** | G4 | None | None | High<sup>1</sup> | High<sup>1</sup> | 83.2 | 83.0 | 80.2 | 78.1 | 75.3 | 74.4 | 72.6 |
|  | Duplex | 73.9 | 77.4 | 73.8 | 66.7 | 48.8 | 45.6 | 37.0 | 32.2 | 27.0 | Low<sup>2</sup> | Low<sup>2</sup> |
| **Duplex-Dark (Fluorescein-G4 and G4Comp-Quencher)/NaClO<sub>4</sub>** | G4 | n/a | n/a | n/a | n/a | n/a | n/a | n/a | n/a | n/a | n/a | n/a |
|  | Duplex | 75.9 | 82.0 | 80.5 | 69.9 | 58.2 | 53.1 | 43.9 | 37.3 | 30.3 | 27.6 | Low<sup>2</sup> |
| **Fluorescein-G4-Quencher/NaClO<sub>4</sub>** | G4 | 65.1 | 82.7 | 88<sup>1</sup> | 92<sup>1</sup> | 88<sup>1</sup> | 86<sup>1</sup> | 81.3 | 77.8 | 75.1 | 74.9 | 73.7 |
|  | Duplex | n/a | n/a | n/a | n/a | n/a | n/a | n/a | n/a | n/a | n/a | n/a |
| **G4-Dark (Fluorescein-G4-Quencher and G4Comp)/LiClO<sub>4</sub><sup>3</sup>** | G4 | None | None | None | None | None | None | n/d<sup>4</sup> | n/d<sup>4</sup> | n/d<sup>4</sup> | n/d<sup>4</sup> | n/d<sup>4</sup> |
|  | Duplex | 76.6 | 81.5 | 80.2 | 69.9 | 57.7 | 46.6 |  |  |  |  |  |
| **Duplex-Dark (Fluorescein-G4 and G4Comp-Quencher)/LiClO<sub>4</sub><sup>3</sup>** | G4 | None | None | None | None | None | None | n/d<sup>4</sup> | n/d<sup>4</sup> | n/d<sup>4</sup> | n/d<sup>4</sup> | n/d<sup>4</sup> |
|  | Duplex | 79.8 | 84.7 | 83.0 | 72.7 | 62.6 | 50.1 |  |  |  |  |  |

<sup>1</sup>Unfolding transition not complete at 95 °C so sigmoid fit not possible, number reported obtained using first derivative method where possible
<sup>2</sup>Folding transition not complete at 20 °C
<sup>3</sup>Only duplex-single stranded transition is observed in this system due to the absence of sodium and reporters used.
<sup>4</sup>Not determined owing to the lower solubility of LiClO<sub>4</sub> vs. NaClO<sub>4</sub>.

We next examined the thermal response of the **Fluorescein-G4-Quencher** component of **G4-Dark** in NaClO_4_-containing solution to ascertain the impact of this salt on the stability of the G-quadruplex states of this system (**Figure 2c**, **Supplementary Figures 7-17**). In all NaClO_4_-containing solutions, **Fluorescein-G4-Quencher** formed a G-quadruplex, giving a 65.1 °C thermal midpoint in 100 mM NaClO_4_; this temperature increased with increasing NaClO_4_, consistent with the electrostatic screening afforded by Na^+^ as well as its binding to the central channel within the G-quadruplex. Between 1-4 M NaClO_4_, the G-quadruplex was so stable it was not fully denatured, even at 95 °C (representative melting curve of 4 M NaClO_4_ in **Figure 2c**). In 5 M NaClO_4_ and above, thermal midpoints were measurable and decreased with increasing perchlorate but remained high. Even in saturated NaClO_4_ (ca. 9 M and containing less than three water molecules per ion), **Fluorescein-G4-Quencher** exhibited a higher thermal midpoint (73.7 °C) than in 100 mM NaClO_4_ (65.1 °C).

The destabilization of the DNA duplex in **G4-Dark** and **Duplex-Dark** was marked (Figure **2i**) and exhibited a near-linear response in thermal midpoint of −7 °C/M NaClO_4_ between 1 and 9 M (r^2^=0.97); these duplexes exhibited thermal midpoints of ca. 75 °C at 100 mM NaClO_4_ and ca. 27-30 °C at 7-8 M NaClO_4_. In contrast, perchlorate-induced destabilization of the G-quadruplex formed by the G-rich strands was less pronounced and/or more than compensated by the presence of additional sodium (**Figure 2d**). **Fluorescein-G4-Quencher** exhibited a thermal midpoint of 65.1 °C at 100 mM NaClO_4_ and 73.7 °C in saturated sodium perchlorate (ca. 9 M). That is, in a polymorphic sequence, one possible secondary structure (dsDNA) that is *more* stable in low perchlorate than an alternative fold (a G-quadruplex) becomes *less* stable than the alternative fold in high perchlorate.

Because of this, we speculated that **G4-Dark** could be thermally switched between all three states (duplex, G-quadruplex, and single-stranded) at intermediate perchlorate concentrations. (**Figure 2e, Supplementary Figures 18-28**). Consistent with this, we observed a single transition at low (0.1 M) NaClO_4_, which corresponded to the duplex-to-single-strand transition. At high NaClO_4_ (saturated/ca. 9 M), we also observed a single transition, which corresponded to the G-quadruplex to single-strand transition (**Figure 2e**), confirmed by the lack of thermal dequenching under these conditions with **Duplex-Dark** (**Figure 2f**).

We next sought to characterize the reversibility of cycling through the system’s states. Reversibility without degradation is essential to a switchable catalyst and a potential concern given some structure-switching nanodevices’ propensity to exhibit degradation with repeated switching due to buildup of waste products.^17,19^ To do so, we thermally cycled a sample of **G4-Dark** in 4 M NaClO_4_ (conditions in which all three states are thermally accessible) 100 times while monitoring fluorescence. The sample did not exhibit degradation during this experiment (**Figure 3a-d)**.

**Figure 3.**
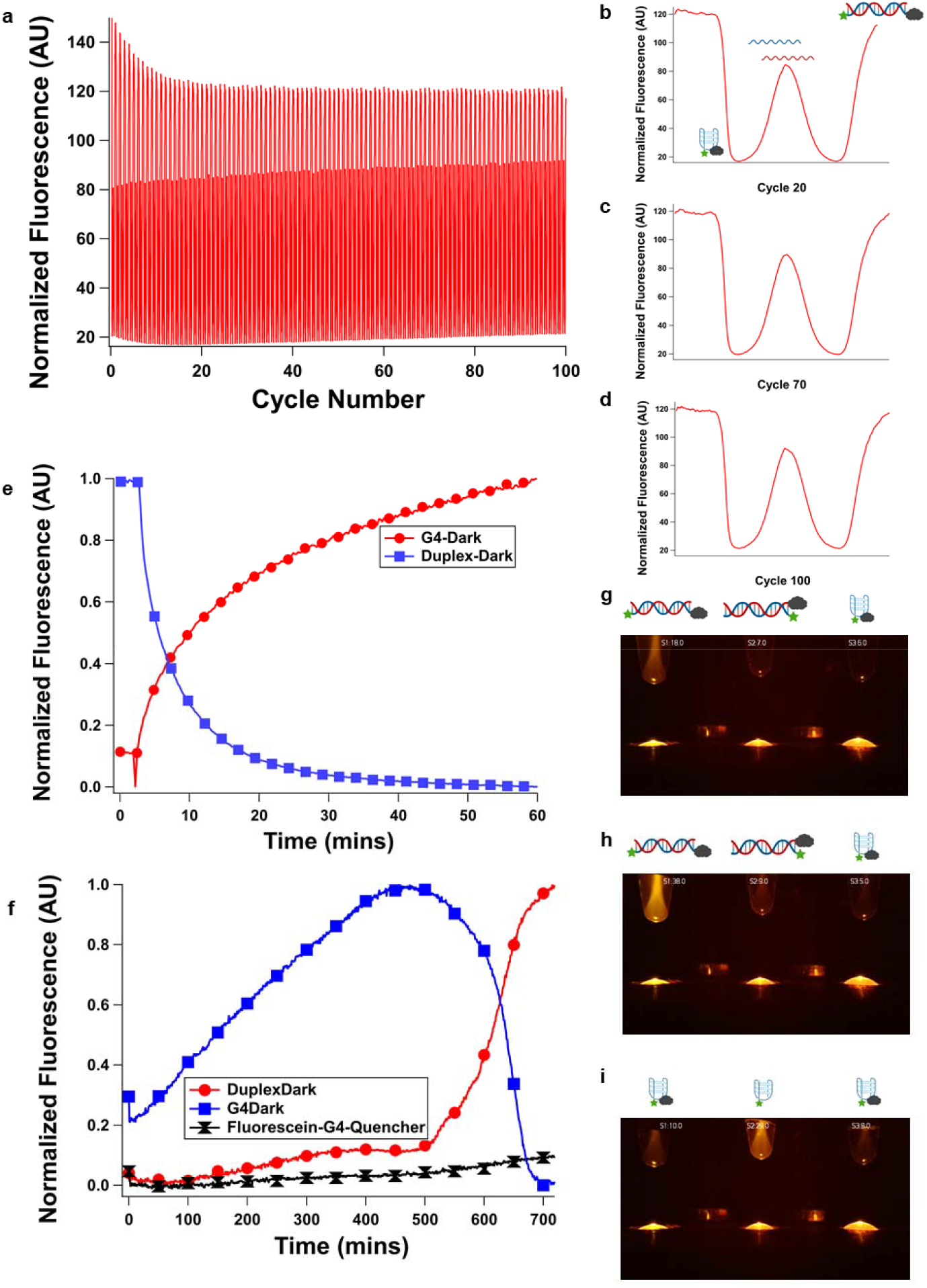
Switching behavior of **G4-Dark**. **G4-Dark** (which reports on all three states) was switched thermally 100 times between duplex, G-quadruplex, and single-stranded states with no decay of fluorescence signal following an initial 10-cycle “burn-in” (panel **a** shows the full series of switching events; panels **b-d** show cycles 20, 70 and 100). While thermal switching is rapid, a “latching” behavior with slower switching is possible with chemical switching, enabling a high level of temporal control of device switching. To demonstrate this, **G4-Dark** and **Duplex-Dark were** also switched chemically by dilution on slower timescales, enabling refolding from the G-quadruplex to duplex state (panel **e**) and vacuum concentration, enabling refolding from the duplex to G-quadruplex state. In vacuum experiments, **G4-Dark** (left) **Duplex-Dark** (center) and **Fluorescein-G4-Quencher** (right) were employed (panel **f** shows fluorescence quantitated from images, panels **g-i** show various timepoints, and this experiment is shown in Supplementary Videos 1 and 2). Under all conditions, **Fluorescein-G4-Quencher** remained a G-quadruplex and did not emit fluorescence. At low (0.8 M) NaClO_4_, the duplex state was favored, resulting in **G4-Dark** emitting fluorescence and **Duplex-Dark** existing in a fluorescence-quenched state (panel **g**). At intermediate perchlorate concentrations during vacuum concentration, the fluorescence of **G4-Dark** increased due to an increase in oligonucleotide concentration but an insufficient increase in NaClO_4_ concentration to induce a structure change; **G4-Dark** remained fluorescent and **Duplex-Dark** remained in a quenched state due to this (panel **h**). Finally, upon ca. 10-fold concentration (to ca. 8 M NaClO_4_, meniscus demonstrating ca. 10-fold concentration from 200 to 20 μL visible) **G4-Dark** and **Duplex-Dark** assumed their G-quadruplex states, causing **G4-Dark** to quench and **Duplex-Dark** to become fluorescent (panel **i**). Brightness was increased by 50% in panels **g-i** relative to captured images for visual clarity. Original images with plots of image-quantitated fluorescence and annotations are shown in Supplementary Videos 1 and 2.

Given the results from **Figure 2**, we reasoned that it would be possible to switch the state of our catalyst from G-quadruplex to duplex (by diluting it with aqueous buffer, lowering the concentration of perchlorate) or from duplex to G-quadruplex (by removing water under vacuum, increasing the concentration of perchlorate). To do so, we performed dilutions of high-salt solutions of **G4-Dark** (which would initially exist in its dark state and transition to its light state) and **Duplex-Dark** (which would initially exist in its light state and transition to its dark state). **G4-Dark** recovered fluorescence upon dilution from 8 M to 0.8 M, and **Duplex-Dark** lost fluorescence following the same dilution (**Figure 3e**). Conversely, we sought to switch the system by removal of solvent. To do so, we took samples with an initial volume of 200 μL and initial concentration of 0.1 M NaClO_4_ and placed them in a vacuum chamber. We monitored these samples by fluorescence imaging with a custom-built device (**Supplementary Figures 29-36**). The samples, initially in low NaClO_4_, behaved as expected for the duplex state, with **G4-Dark** exhibiting fluorescence and **Duplex-Dark** in a dark state. **Duplex-Dark** remained dark during concentration while the fluorescence of **G4-Dark** gradually increased as the solution became more concentrated. Finally, both solutions reached a critical concentration of NaClO_4_ at which they transitioned to the G-quadruplex state: **Duplex-Dark** became fluorescent and **G4-Dark** became nonfluorescent (**Figure 3f-i, Supplementary Videos 1 and 2, Supplementary Code 1-3**).

**G4-Dark** and **Duplex-Dark** demonstrate the perchlorate-based switchability of our system. To demonstrate the ability of G-quadruplex/duplex equilibria to enable switchable electron transfer, we constructed **G4-SwitchR** (switchable redox; **Table 1**), which was comprised of **G4Redox**, **G4Redox-Comp**, and the cofactor hemin. We employed Amplex Red, a nonfluorescent dye that is oxidized to the fluorescent, red-colored pigment resorufin catalytically by the peroxidase-mimicking DNAzyme formed between a G-quadruplex and hemin (**Figure 4a-c**).^26^

**Figure 4.**
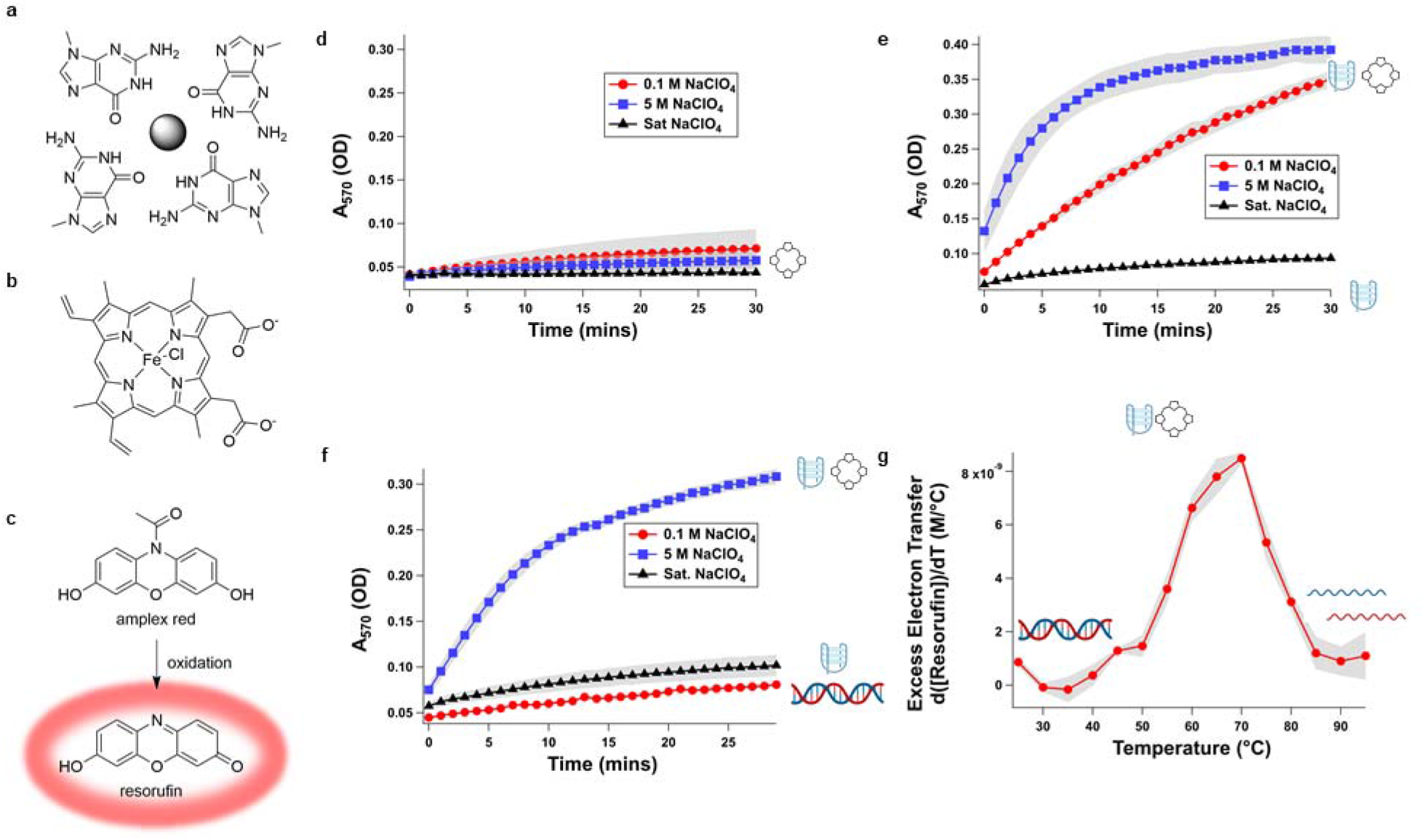
Switchable electron transfer amplification behavior of **G4-SwitchR** and its constituent components in the presence of hemin. When folded into the G-quadruplex state, a solvent-exposed G-quartet (panel **a**) is present which is known to provide a tight binding pocket for the cofactor hemin (panel **b**); the complex formed by a G-quadruplex and hemin can catalyze electron transfer. We visualize this here using the colorimetric electron transfer probe Amplex Red (panel **c,** top), which is initially colorless and nonfluorescent and, when oxidized, converts to the red fluorescent pigment resorufin (panel **c**, bottom). Hemin itself has a minimal background rate of electron transfer that is suppressed by increasing NaClO_4_ concentration (panel **d**). When **G4-Redox** is operated as the only oligonucleotide constituent (i.e., under conditions in which the strand will form only a G-quadruplex) in the presence of hemin, robust multiple-turnover electron transfer occurs in low (0.1 M, red circles) NaClO_4_ solution, it is enhanced in moderate (5 M, blue squares) NaClO_4_ solution, and it is suppressed in saturated NaClO_4_ solution (black triangles) due to abrogation of hemin binding (panel **e**). When **G4-SwitchR** is operated in these same conditions, electron transfer is switched off in low (0.1 M) NaClO_4_ solution due to duplex formation, switched on in moderate (5 M) NaClO_4_ solution due to G-quadruplex formation with concomitant hemin binding, and switched off in saturated NaClO_4_ solution due to abrogation of hemin binding (panel **f**). **G4-SwitchR** can also be switched thermally, exhibiting varying rate of reaction depending on its folding state. When ramped from 25 to 95 °C in 5 M NaClO_4_ solution, excess electron transfer (as measured by rate of resorufin production in excess of that observed in hemin alone) starts at near-zero as the device exists in duplex form, increases substantially between 50 and 70 °C as the duplex melts and the G-quadruplex forms, and it decreases again by ca. 90% between 70 and 90 °C as the G-quadruplex unfolds and the single-stranded form of the device forms (panel **g**). These results are consistent with UV-visible monitored thermal measurements of secondary structures formed (**Supplementary Figures 38** and **39**). Error bar shading=S.E.M., n=3 technical replicates.

The potential difference in this system is defined by the concentration of terminal electron acceptor (hydrogen peroxide) and the redox chemistry of the probe. These are held constant between experiments, we compared the ratio of the rate of electron transfer performed by the G-quadruplex/hemin complex to that performed by hemin alone (**Figure 4d**). In 0.1 and 5 M NaClO_4_, the **G4-Redox** component of **G4-SwitchR** binds hemin and can perform electron transfer, with enhanced activity in 5 M relative to 0.1 M NaClO_4_ (**Figure 4e, Supplementary Table 1, Supplementary Figures 37**). In saturated NaClO_4_, electron transfer is suppressed due to lack of hemin-G-quadruplex interactions.

In 0.1 M NaClO_4_, **G4-SwitchR** exists entirely as a duplex, hemin binding is thus abrogated; resorufin was produced at < 30 nM/min when **G4-SwitchR** was present or when it was absent **(Figure 4f)**. In 5 M NaClO_4_, **G4-SwitchR** dissociates into **G4Redox** folded into a G-quadruplex, which is competent to bind hemin, and **G4Redox-Comp** exists as a single strand. The resulting hemin**-G4Redox** complex performs electron transfer (as measured by the reporter dye Amplex Red’s conversion to resorufin) 35-fold more rapidly than hemin alone (**Supplementary Figures 37**): the hemin-**G4Redox** complex produced resorufin at 300 nM/min vs. unbound hemin, which produced it at 9 nM/min. At still higher NaClO_4_, hemin-G-quadruplex binding decreases, and in saturated (ca. 9 M) NaClO_4_, resorufin was produced at a much lower rate of 46 nM/min. Thus, perchlorate concentration can be varied to switch the state of the catalyst as well as to modulate the reaction rate.

In order to demonstrate electron transfer could be thermally switched as well, we measured Amplex Red oxidation using **G4-SwitchR** in 4 M NaClO_4_ (**Figure 4g**). At 20 °C, electron transfer was near-zero. As the sample was heated and switched from duplex to G-quadruplex, the rate of electron transfer exhibited a significant increase, reaching a maximum at 60 °C and again decreasing as the G-quadruplex unfolded. This is consistent with the thermal denaturation curve generated by monitoring A_295_ (a diagnostic wavelength for G-quadruplex formation) of **G4-SwitchR** (**Supplementary Table 1**, **Supplementary Figures 38-39**). Notably, hydrogen peroxide was required. Perchlorate ion, despite its exceptionally high oxidation potential, did *not* suffice as an electron acceptor in this system, demonstrating the exceptional specificity afforded by biocatalyst-based electronic systems and the compatibility of this high-oxidation state ion-that is ubiquitous in legacy electronic systems-with biomolecular electronics.

## Discussion, Conclusions, and Outlook

Here, we have demonstrated that DNA-hemin complexes can act as electron transfer catalysts that can be switched by exploiting the phenomenon we have observed in which duplex DNA is preferentially destabilized by perchlorate salts relative to G-quadruplex DNA. This phenomenon can be used to switch electron transfer behavior in three ways: 1) direct addition of perchlorate salts, 2) thermal switching, and 3) increasing perchlorate salt concentration by vacuum concentration.

The electron transfer phenomena shown here suggest the possibility exists to interface such biopolymers with conventional electronics, as was recognized as a Key Technical Area by the Semiconductor Research Corporation in their Biocomputing Roadmap.^27^ Additionally, G-quadruplex catalysts are used for many applications such as use as an oxidizing agent for disulfide bonds,^28^ self-biotinylation,^29^, and Diels-Alder transformations,^30^ This suggests that the possibility exists to use switchable G-quadruplexes for a wide range of conditional transformations. Th

Examples of secondary-structure remodeling are observed in life as well. Nucleic acids exhibit considerable polymorphism in biological systems, particularly in sequences that can form G-quadruplexes. Such systems are diverse and widespread in biology and extend to both DNA and RNA, including proto-oncogene promoters,^31^ the expansion segments of rRNA of higher organisms,^32–34^ the rDNA corresponding to those expansion segments,^35^ and eukaryotic messenger RNAs.^13,14^ Several proteins can remodel G-quadruplexes, and both energy-dependent (i.e., helicases) and energy-independent systems that can do so have been reported.^36,37^ ATP-dependent helicases are known to unwind G-quadruplex structures, and the RNA-binding protein Lin28 has been shown to unfold G-quadruplexes without the requirement for ATP.^36^ The structure switching performed by Hofmeister ions in this work amounts to a bioinspired means of performing such remodeling.

Given that such remodeling processes are also operative in nature and that hemin-G-quadruplex promoted electron transfer has been suggested as being physiologically relevant, we speculate that conditional hemin-G-quadruplex complexes are a means by which cells could conditionally enable electron transfer.^38^ For example, ribosomes from the neurons of Alzheimer’s Disease patients have been shown to contain more iron than those from healthy patients, and these ribosomes possess peroxidase activity.^39^ G-quadruplexes may enable this phenomenon *in vivo*. Human ribosomes are known to be polymorphic, particularly in their G-rich expansion segments, some of which have been observed to lack electron density in EM maps, consistent with an equilibrium between multiple states.^32,40^ Rarely, individual ribosomes clearly exhibit extended conformations consistent with Watson-Crick base pairing, but these same sequences possess exceptionally high G-quadruplex forming potential and form G-quadruplexes in cell-free *in vitro* experiments, which is consistent with polymorphism with one state capable of catalyzing electron transfer.^34,35^

We suggest that conditionally folded G-quadruplexes in cells could collaborate with hemin, producing a means by which cells can perform conditional electron transfer that is analogous to the phenomenon we have employed in this work. In fact, recent work suggests that hemin-G-quadruplex associations occur in human cells.^38^ Such a phenomenon could be exploited both by extant life or in synthetic biological systems, and we speculate this could have enabled conditionally active forms of prebiotic electron transfer catalysts, which have attracted intense interest in recent years.^6,41,42^

## Acknowledgements

Several figures in this manuscript were made using BioRender.com. We thank Loren Williams and George Perry for helpful discussions. The imaging apparatus used here was developed as part of the RockSat-C 2018 sounding rocket program, sponsored by the Colorado Space Grant Consortium and NASA Wallops; we thank the NASA PAXC team members for helpful discussions. This work was supported by NASA Contract 80NSSC18K1139 under the Center for Origin of Life (to A.E.E. and K.P.A.).

## Data Availability Statement

The datasets generated during and/or analyzed during the current study are available from the corresponding author on reasonable request.

## Code Availability Statement

All computer code used during the current study is included in this published article (and its supplementary information files).

## Author Contributions

A.E.E., K.P.A., T.G.H., and M.R.P. conceived the project. T.G.H., M.R.P., L.M.A., and B.R.B. performed experiments. L.M.A. contributed computer codes and analysis. M.B.G. performed CAD of the fluorescence imaging jig. N.B.B. fabricated and assembled the fluorescence imaging jig. A.E.E., K.P.A., T.G.H., and M.R.P. wrote the paper.

## Methods

### Nucleic Acids

Oligonucleotides were obtained from Integrated DNA Technologies (Coralville, IA) and used as received. Labeled strands were obtained with HPLC purification and unlabeled strands were obtained with standard desalting.

### DNA Sample Preparation

Oligonucleotides were annealed in a T100 thermal cycler (Bio-Rad) by incubating at 95 °C for 2 minutes, then decreasing the temperature by 10 °C steps and annealing at each step for 1 minute to a final temperature of 25 °C. Except where otherwise specified, experiments were performed in 50 mM Li-HEPES, pH 7.4, with 1 μM of the duplexed or single stranded oligonucleotides.

### Fluorescence-Monitored Experiments

Fluorescence measurements were performed using a Gemini XS plate reader (Molecular Devices) or a Cary Eclipse (Varian Technologies) fluorometer equipped with a thermostated peltier holder (Agilent Technologies) when bidirectional temperature control was required. Excitation was performed at 495 nm and emission was monitored at 520 nm.

For thermal denaturation studies, the sample was ramped from 20 °C to 95 °C (two heat/cool cycles) at 5 °C/min. Selected measurements were also performed with a slower 0.5 °C/min ramp rate to mitigate hysteresis. Thermal midpoints are reported as a sigmoid fit of the transition(s) in the second heating trace of two temperature ramps.

Data from fluorescence melts were normalized by division, setting the highest RFU value to 1 and the lowest value to 0 for each state in a given system (quenched or dequenched/partially dequenched).

### UV-Vis Monitored Experiments

Thermal denaturation experiments were performed using a qCHANGER 6/Cary60 (Quantum Northwest) interfaced to a Cary 60 UV-Visible spectrophotometer (Agilent Technologies) using a custom ADL script developed by Quantum Northwest to collect full spectra at each temperature as described previously. Heat sinking for the Peltier device was provided by an EXT-440CU ambient liquid cooling system (Koolance). Duplex melting transitions were monitored using the 260 nm trace from these datasets and G-quadruplex melting temperatures were monitored using the 295 nm trace.

Amplex Red experiments (described further below) were monitored on a Cary 60 UV-vis (Agilent Technologies) or a SpectraMAX 340PC plate reader (Molecular Devices) with the PathCheck functionality enabled.

### Fluorescence Imaging Under Vacuum

Samples were pipetted into PCR tubes with the lids open, which were inserted into a 3D-printed jig that was placed inside a black box. The jig was fabricated in PLA on a Prusa 3D printer. The box and its contents were placed in a vacuum desiccator (Supplementary Figures 29-36) and the chamber was continuously evacuated with a diaphragm pump (Welch 2014B-01). The samples were excited with LEDs with emission centered at 462-465 nm (Item B01GDO9UNY, Amazon). The samples were imaged with a camera (Raspberry Pi Module V2, Amazon) fixed at 90° relative to the light sources. A 12.5 mm longpass filter (Schott OG 550, Edmund Optics) was affixed with polyvinyl acetate adhesive (Elmer’s Glue-All, Costco Wholesale) directly in front of the camera lens to block excitation light and pass emitted light. The camera was interfaced to a Raspberry Pi B+ (Amazon). A Python script controlled the Raspberry Pi’s GPIO pins to illuminate samples at 1-minute intervals, collect images, and measured and plotted fluorescence intensity vs. time. Time-lapse movies were generated from captured images using FFmpeg.

Samples were prepared with a starting volume of 200 μL and 0.1 μM of **G4-Dark, Duplex-Dark,** or **Fluorescein-G4-Quencher** in 5 mM Li-HEPES, pH 7.4 and 0.8 M NaClO_4_. After desiccation, the final volume was 20 μL with 50 mM Li-HEPES, 8 M NaClO_4_ and 1 μM **G4-Dark, Duplex-Dark,** or **Fluorescein-G4-Quencher.**.

### Electron Transfer Assay

Samples contained **G4-SwitchR** (i.e., **G4Redox** and **G4Redox-Comp**), **G4Redox** alone, or no oligonucleotide. **G4Redox**, when present, was at 1 μM and **G4Comp**, when present, was at 2 μM to ensure full duplex state at low perchlorate/temperature and that electron transfer catalysis observed was due to the G-quadruplex state of **G4-SwitchR** and not residual unduplexed **G4Redox**. The buffer used here was 5 mM sodium phosphate, pH 7.4 with varying sodium perchlorate concentrations. To initiate the reaction, hemin was added to a final concentration of 1 μM, hydrogen peroxide was added to a final concentration of 300 μM, and Amplex Red was added to a final concentration of 200 μM. Reactions were monitored by measuring the absorbance of resorufin at 570 nm. For the Cary 60 thermal activation-Amplex red assay, Amplex Red was added to a final concentration of 2 mM. All reactions contained 2% DMSO, which came from the hemin and Amplex Red stocks

