## Supplementary Materials for "Minimal DNA Electron Transfer Catalysts Switched by a Chaotropic Ion"

for

**Table of Contents**

Supplementary Figure 1. Fluorescence melting curve (second heating trace) of **Duplex-Dark** in 0.1 M LiClO_4_.

Supplementary Figure 2. Fluorescence melting curve (second heating trace) of **Duplex-Dark** in 0.5 M LiClO_4_.

Supplementary Figure 3. Fluorescence melting curve (second heating trace) of **Duplex-Dark** in 1 M LiClO_4_.

Supplementary Figure 4. Fluorescence melting curve (second heating trace) of **Duplex-Dark** in 2 M LiClO_4_.

Supplementary Figure 5. Fluorescence melting curve (second heating trace) of **Duplex-Dark** in 3 M LiClO_4_.

Supplementary Figure 6. Fluorescence melting curve (second heating trace) of **Duplex-Dark** in 4 M LiClO_4_.

Supplementary Figure 7. Fluorescence melting curve (second heating trace) of **Fluorescein-G4-Quencher** in 0.1 M NaClO_4_.

Supplementary Figure 8. Fluorescence melting curve (second heating trace) of **Fluorescein-G4-Quencher** in 0.5 M NaClO_4_.

Supplementary Figure 9. Fluorescence melting curve (second heating trace) of **Fluorescein-G4-Quencher** in 1 M NaClO_4_.

Supplementary Figure 10. Fluorescence melting curve (second heating trace) of **Fluorescein-G4-Quencher** in 2 M NaClO_4_.

Supplementary Figure 11. Fluorescence melting curve (second heating trace) of **Fluorescein-G4-Quencher** in 3 M NaClO_4_.

Supplementary Figure 12. Fluorescence melting curve (second heating trace) of **Fluorescein-G4-Quencher** in 4 M NaClO_4_.

Supplementary Figure 13. Fluorescence melting curve (second heating trace) of **Fluorescein-G4-Quencher** in 5 M NaClO_4_.

Supplementary Figure 14. Fluorescence melting curve (second heating trace) of **Fluorescein-G4-Quencher** in 6 M NaClO_4_.

Supplementary Figure 15. Fluorescence melting curve (second heating trace) of **Fluorescein-G4-Quencher** in 7 M NaClO_4_.

Supplementary Figure 16. Fluorescence melting curve (second heating trace) of **Fluorescein-G4-Quencher** in 8 M NaClO_4_.

Supplementary Figure 17. Fluorescence melting curve (second heating trace) of **Fluorescein-G4-Quencher** in Saturated NaClO_4_.

Supplementary Figure 18. Fluorescence melting curve (second heating trace) of **G4-Dark** in 0.1 M NaClO_4_.

Supplementary Figure 19. Fluorescence melting curve (second heating trace) of **G4-Dark** in 0.5 M NaClO_4_.

Supplementary Figure 20. Fluorescence melting curve (second heating trace) of **G4-Dark** in 1 M NaClO_4_.

Supplementary Figure 21. Fluorescence melting curve (second heating trace) of **G4-Dark** in 2 M NaClO_4_.

Supplementary Figure 22. Fluorescence melting curve (second heating trace) of **G4-Dark** in 3 M NaClO_4_.

Supplementary Figure 23. Fluorescence melting curve (second heating trace) of **G4-Dark** in 4 M NaClO_4_.

Supplementary Figure 24. Fluorescence melting curve (second heating trace) of **G4-Dark** in 5 M NaClO_4_.

Supplementary Figure 25. Fluorescence melting curve (second heating trace) of **G4-Dark** in 6 M NaClO_4_.

Supplementary Figure 26. Fluorescence melting curve (second heating trace) of **G4-Dark** in 7 M NaClO_4_.

Supplementary Figure 27. Fluorescence melting curve (second heating trace) of **G4-Dark** in 8 M NaClO_4_.

Supplementary Figure 28. Fluorescence melting curve (second heating trace) of **G4-Dark** in Saturated NaClO_4_.

Supplementary Figure 29. Engineering drawing of 3D printed tube holder/camera mount component of imaging jig. Dimensions in mm.

Supplementary Figure 30. Engineering drawing of 3D printed led holder component of imaging jig. Dimensions in mm.

Supplementary Figure 31. CAD Mockup of Fluorescence Imaging Jig

Supplementary Figure 32. CAD Mockup of Fluorescence Imaging Jig (Alternate view)

Supplementary Figure 33. Imaging jig with tubes.

Supplementary Figure 34. Imaging jig (top view).

Supplementary Figure 35. Imaging Jig (side view)

Supplementary Figure 36. Imaging Jig (front view).

Supplementary Video 1. Fluorescence Imaging shows G4-Dark and Duplex-Dark switching structure under vacuum control with real-time plotting of fluorescence intensity

Supplementary Video 2. Fluorescence Imaging shows G4-Dark and Duplex-Dark switching structure under vacuum control (With annotations and captions)

Supplementary Code 1. Python code for controlling Raspberry Pi for reaction monitoring and overlaying plots of fluorescence vs. time.

Supplementary Code 2. Python code for reanalyzing collected images.

Supplementary Code 3. Python code for generating fluorescence plot overlays for existing image datasets.

Supplementary Table 1. Thermal midpoints of **G4Redox.** Measurements were obtained by UV-vis monitoring of A­_260_ and A­_295_.

Supplementary Figure 37. Kinetic measurements of oxidation of amplex red by **G4Redox** in the presence of varying concentrations of NaClO_4_.

Supplementary Figure 38. Kinetic measurements of oxidation of amplex red by free hemin in the presence of varying concentrations of NaClO_4_.

Supplementary Figure 39. Kinetic measurements of oxidation of amplex red by **G4-SwitchR** in the presence of varying concentrations of NaClO_4_.

Supplementary Figure 40. Endpoint measurements of Amplex Red oxidation by **G4-SwitchR** (Duplexed GQA), **G4Redox** (GQA), and free hemin after 10 minutes.

Supplementary Figure 41. A_295_-monitored melting curve of the **G4-SwitchR** system.

Supplementary Figure 42. A_260_-monitored melting curve of the **G4-SwitchR** system.

Supplementary Figure 1. Fluorescence melting curve (second heating trace) of **Duplex-Dark** in 0.1 M LiClO_4_.

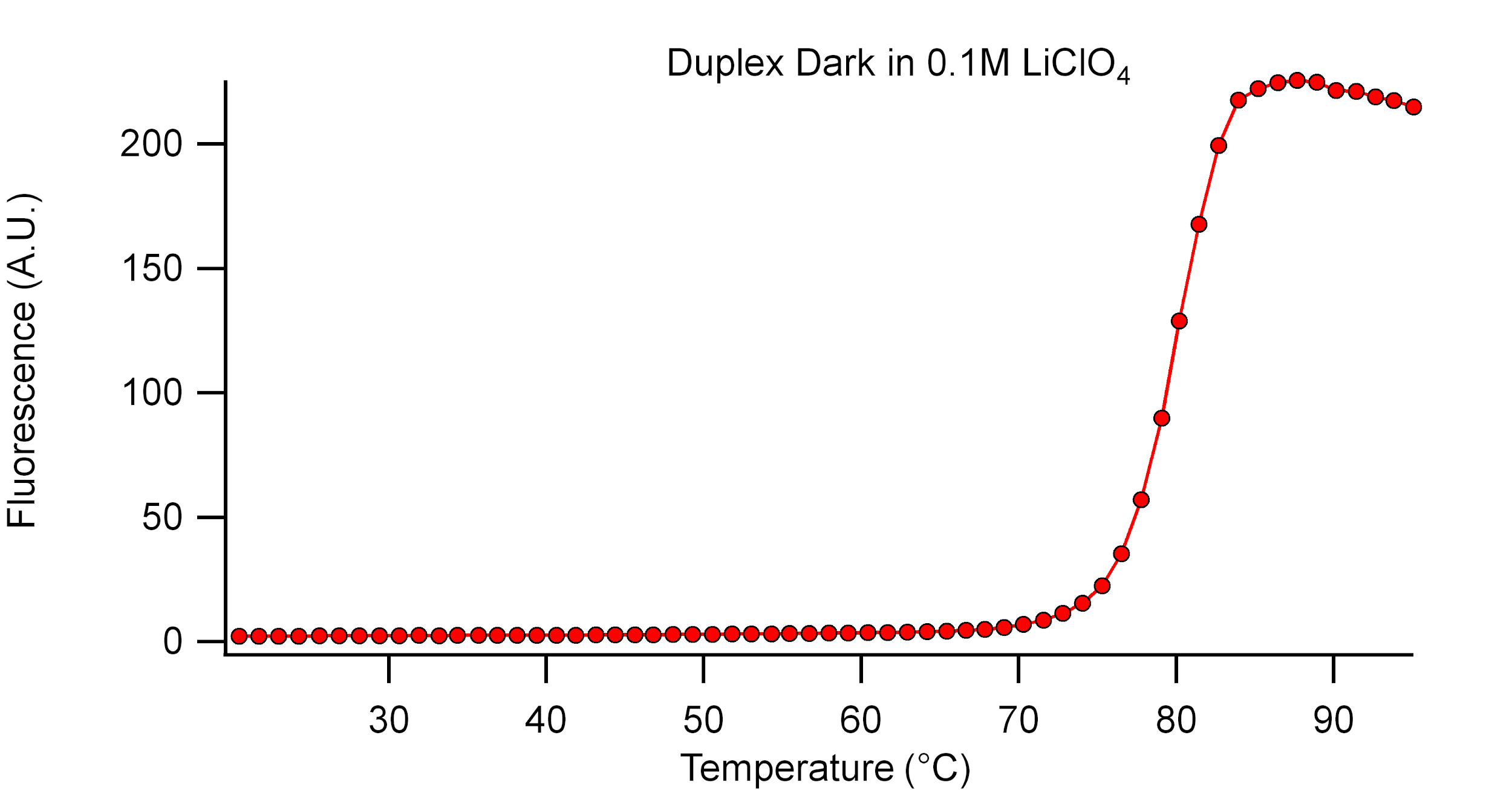

Supplementary Figure 2. Fluorescence melting curve (second heating trace) of **Duplex-Dark** in 0.5 M LiClO_4_.

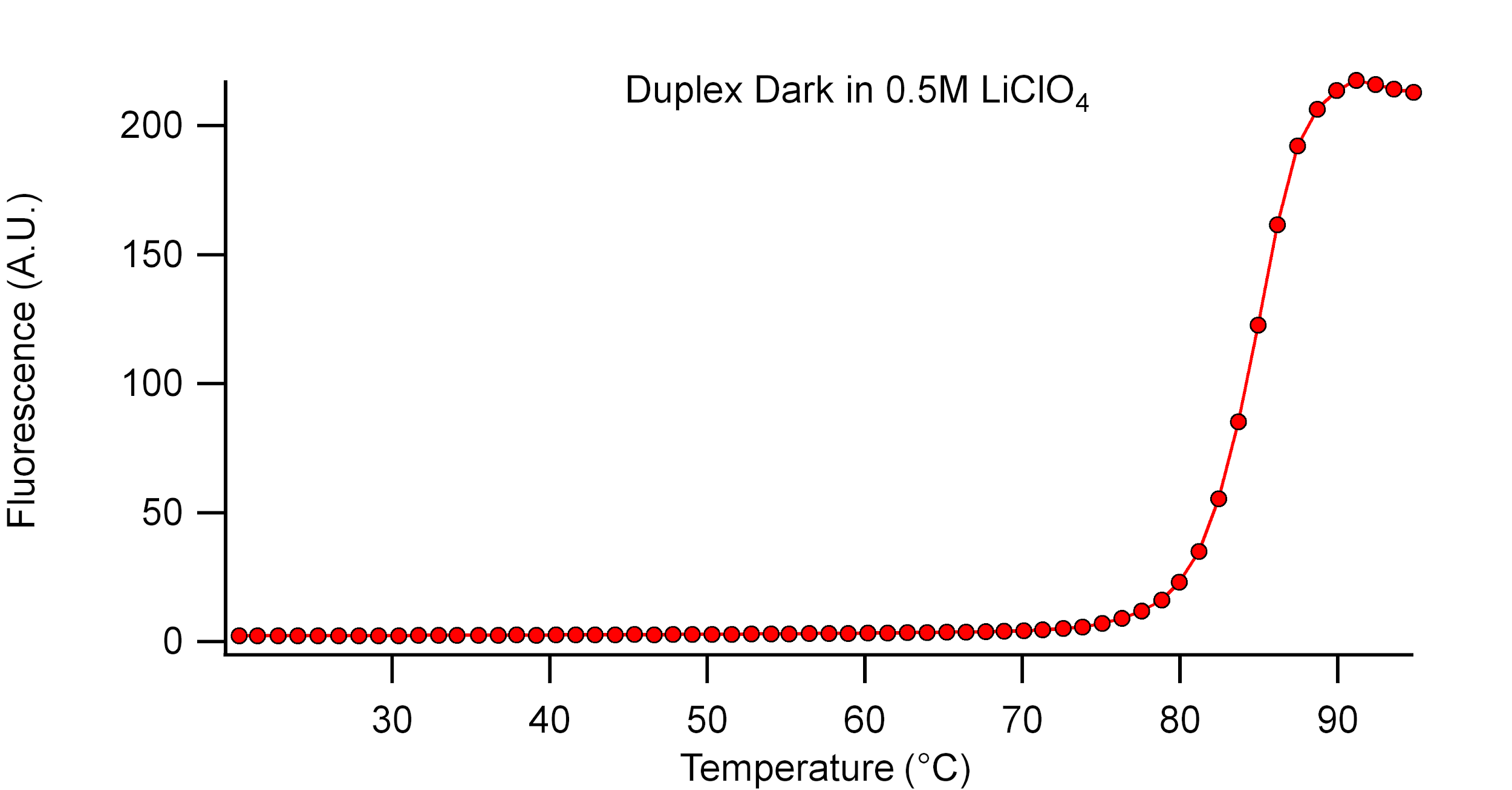

Supplementary Figure 3. Fluorescence melting curve (second heating trace) of **Duplex-Dark** in 1 M LiClO_4_.

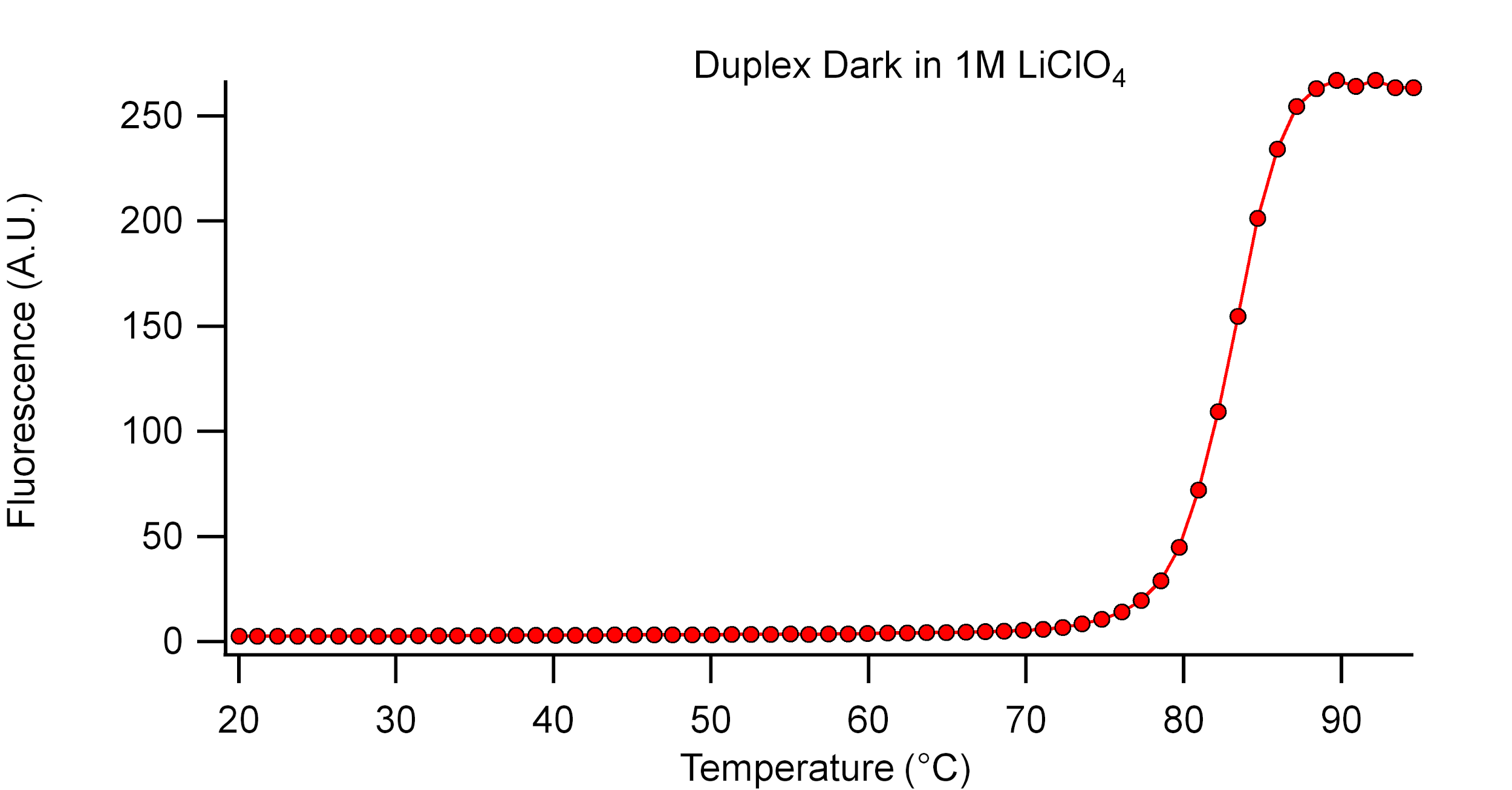

Supplementary Figure 4. Fluorescence melting curve (second heating trace) of **Duplex-Dark** in 2 M LiClO_4_.

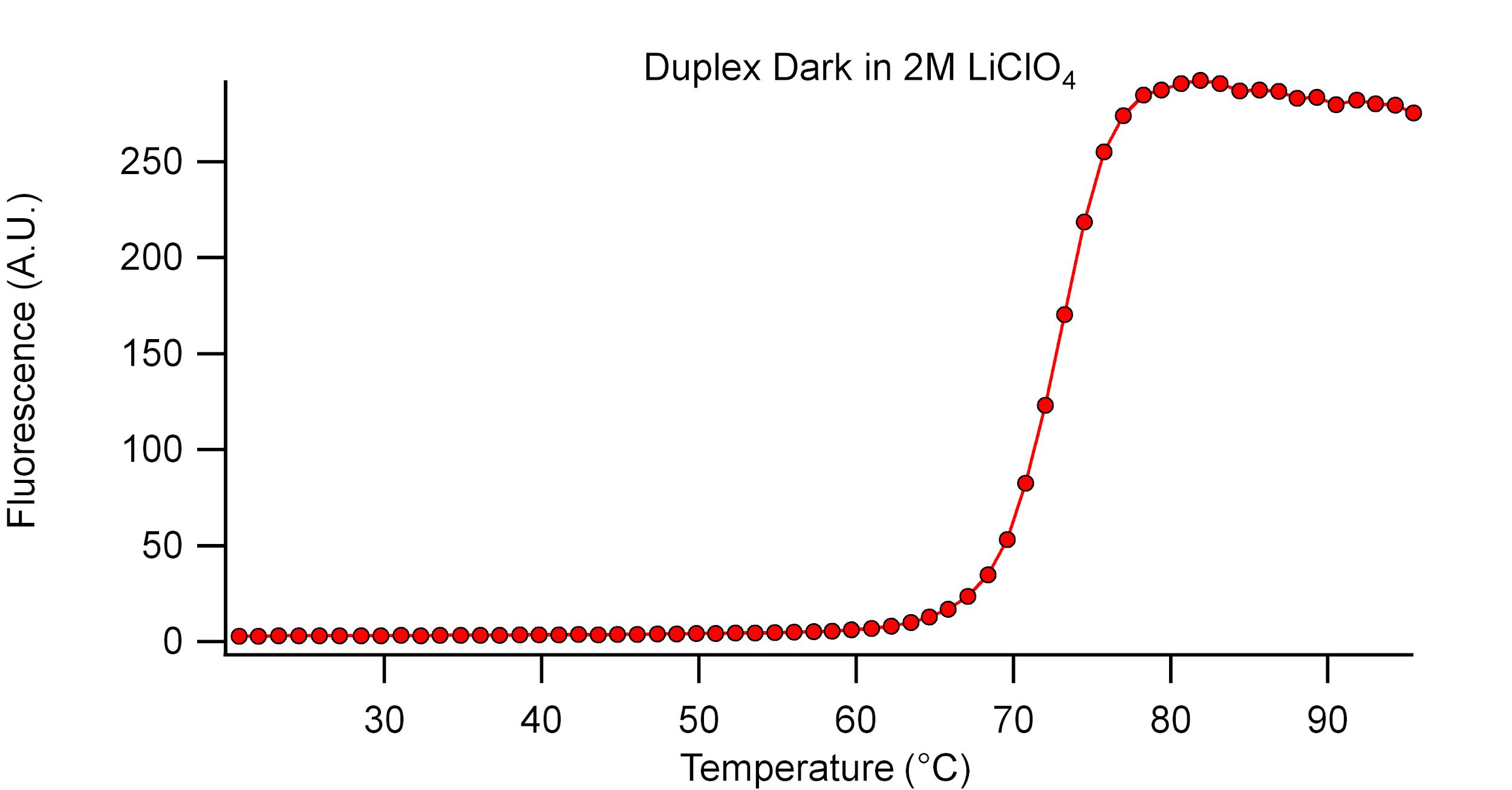

Supplementary Figure 5. Fluorescence melting curve (second heating trace) of **Duplex-Dark** in 3 M LiClO_4_.

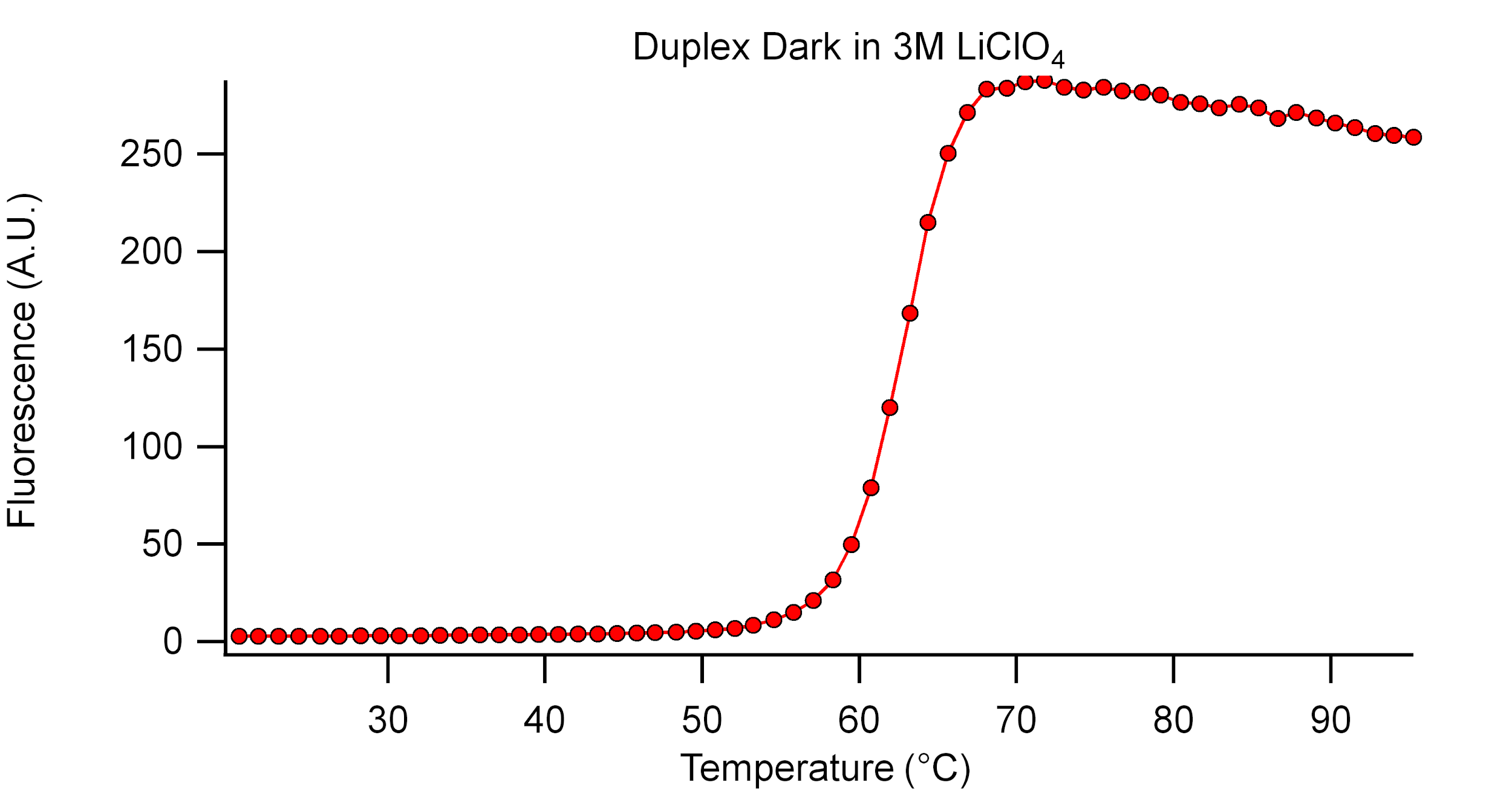

Supplementary Figure 6. Fluorescence melting curve (second heating trace) of **Duplex-Dark** in 4 M LiClO_4_.

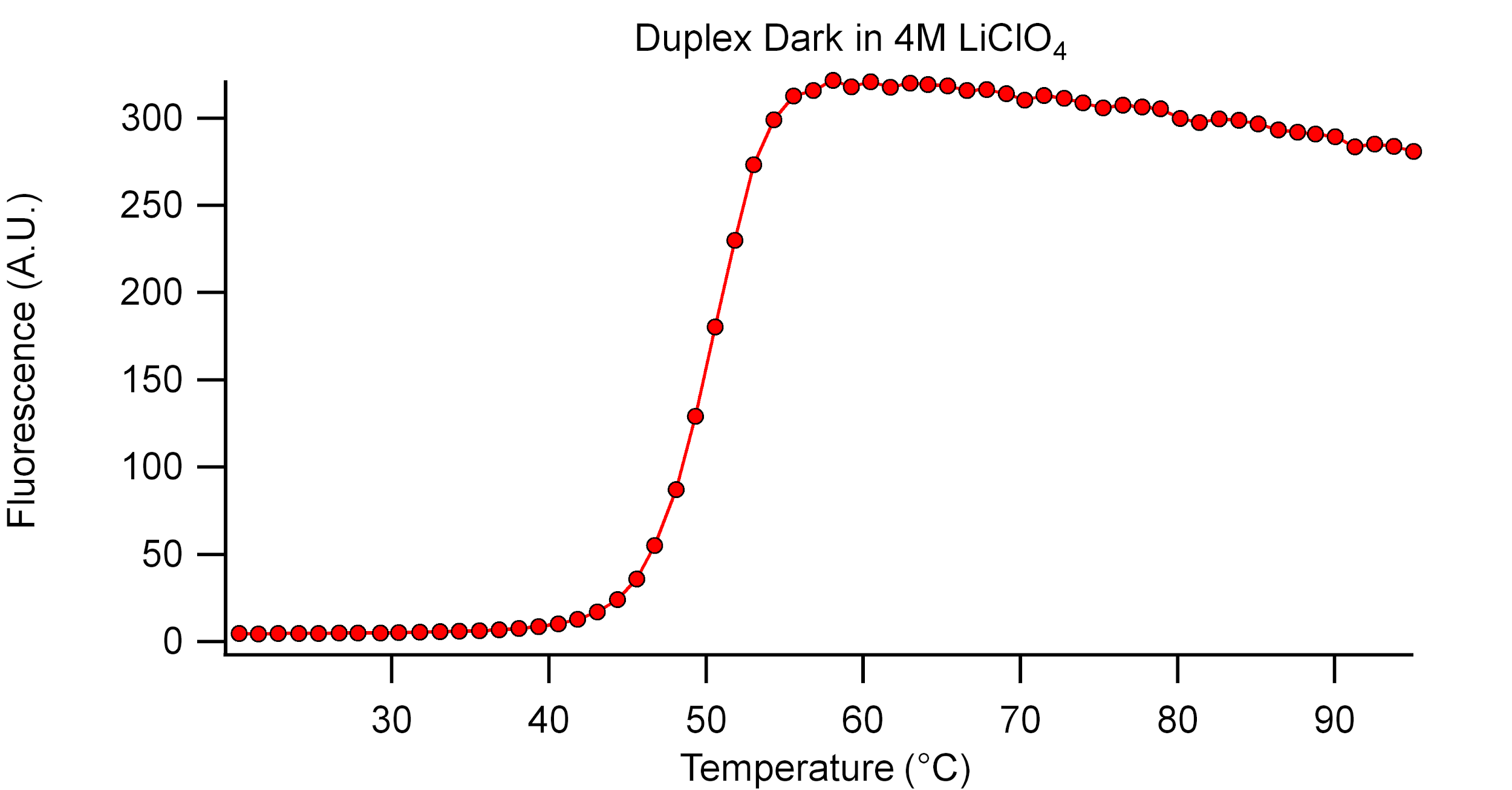

Supplementary Figure 7. Fluorescence melting curve (second heating trace) of **Fluorescein-G4-Quencher** in 0.1 M NaClO_4_.

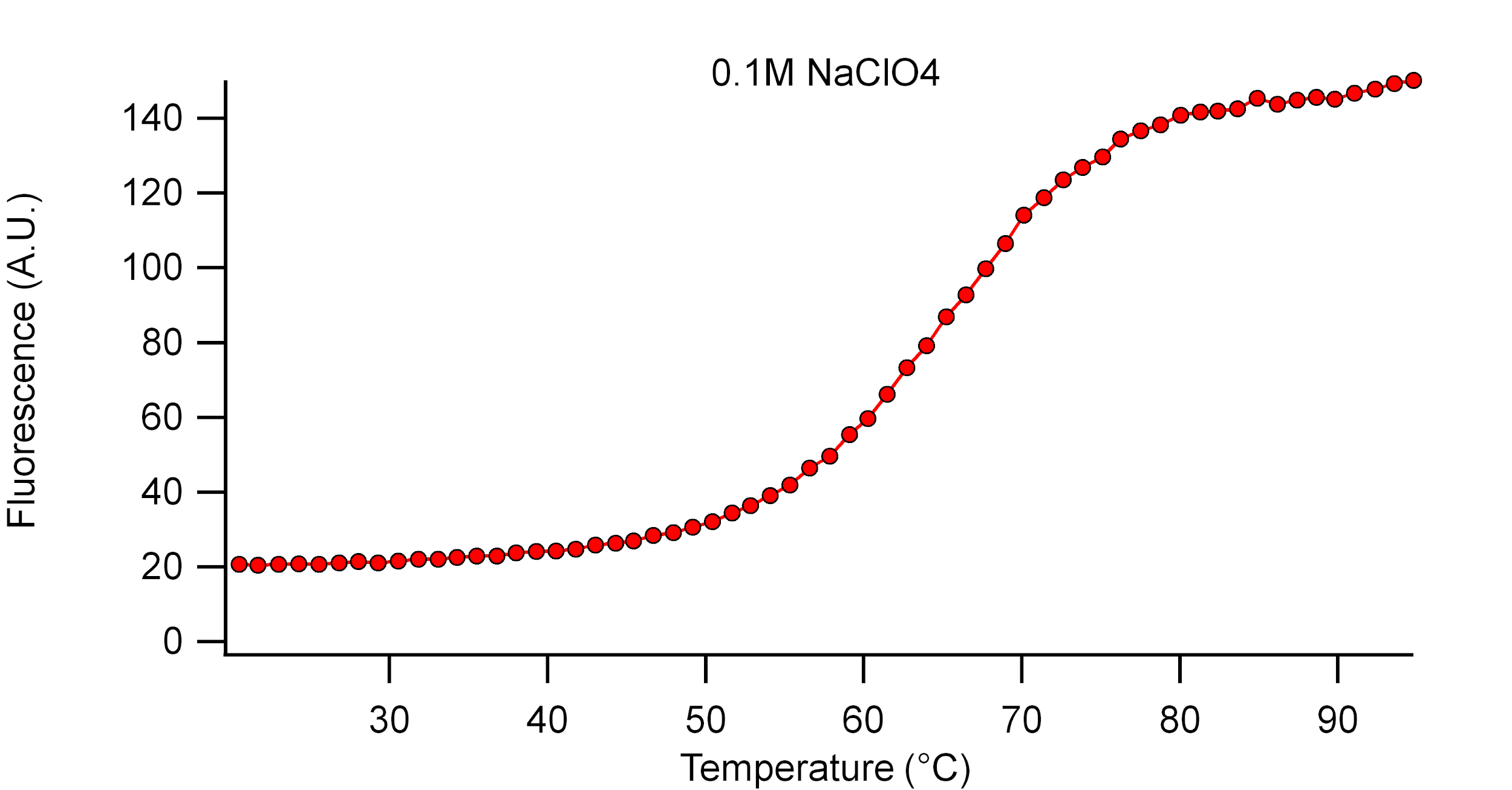

Supplementary Figure 8. Fluorescence melting curve (second heating trace) of **Fluorescein-G4-Quencher** in 0.5 M NaClO_4_.

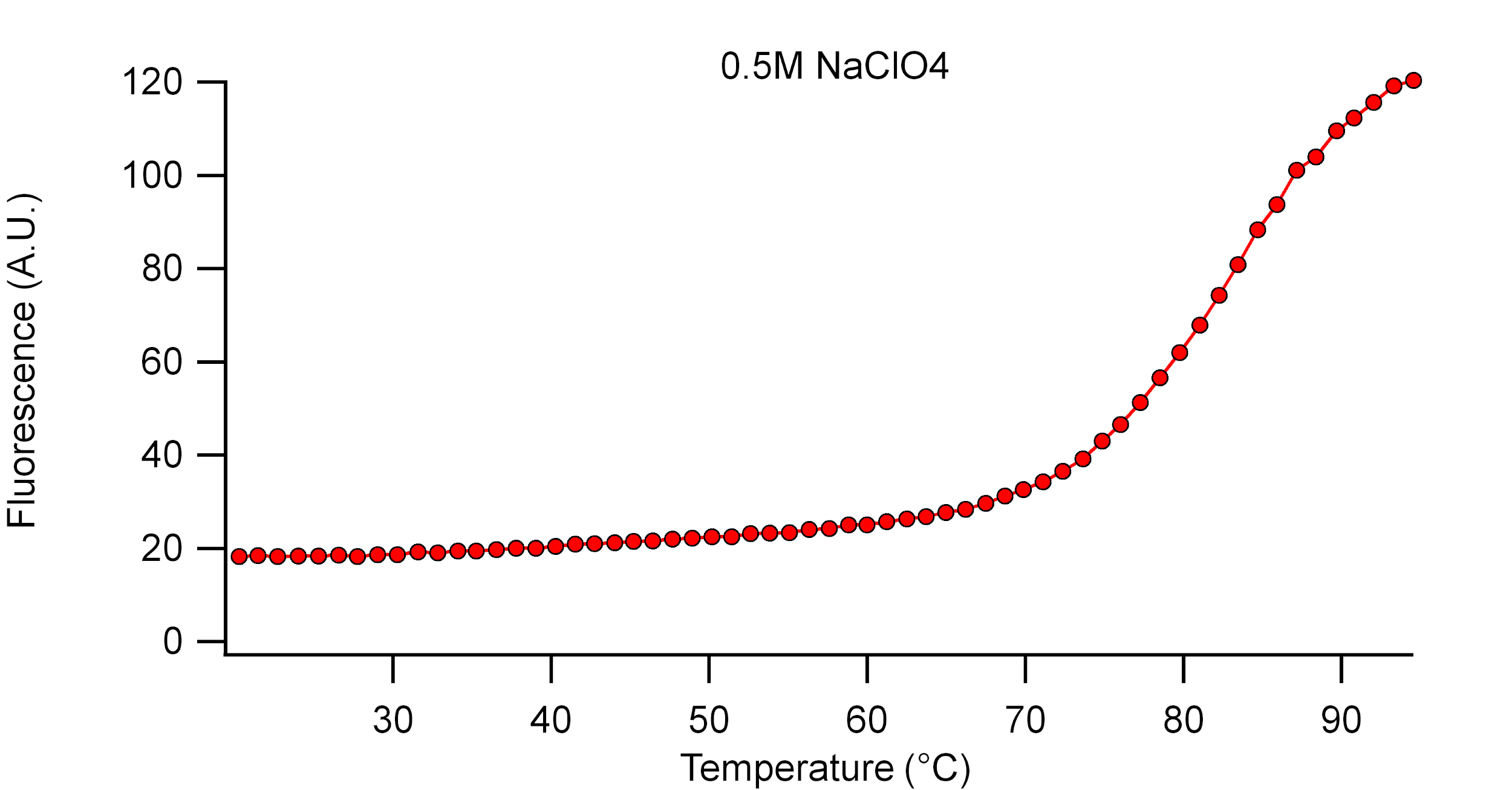

Supplementary Figure 9. Fluorescence melting curve (second heating trace) of **Fluorescein-G4-Quencher** in 1 M NaClO_4_.

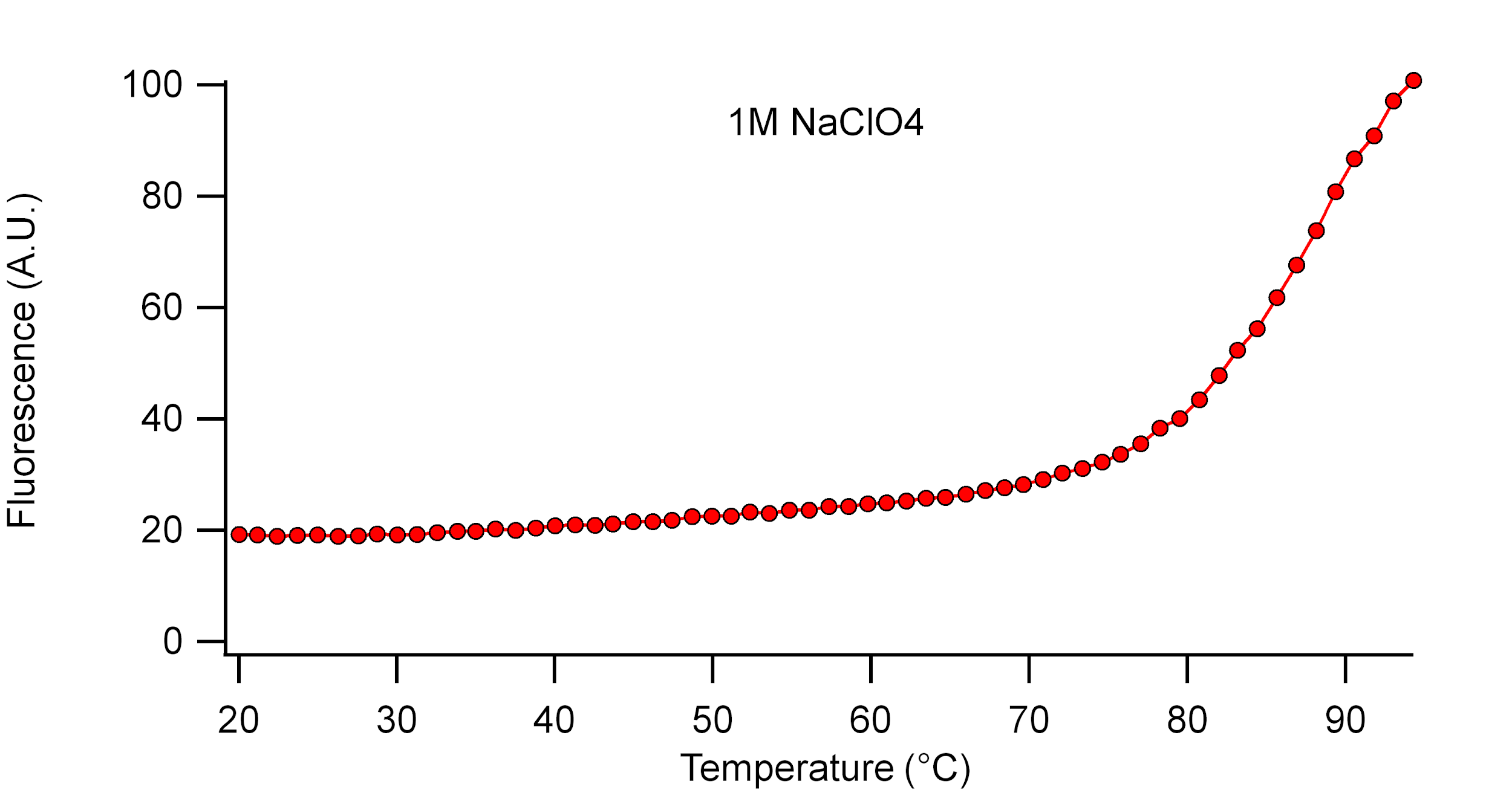

Supplementary Figure 10. Fluorescence melting curve (second heating trace) of **Fluorescein-G4-Quencher** in 2 M NaClO_4_.

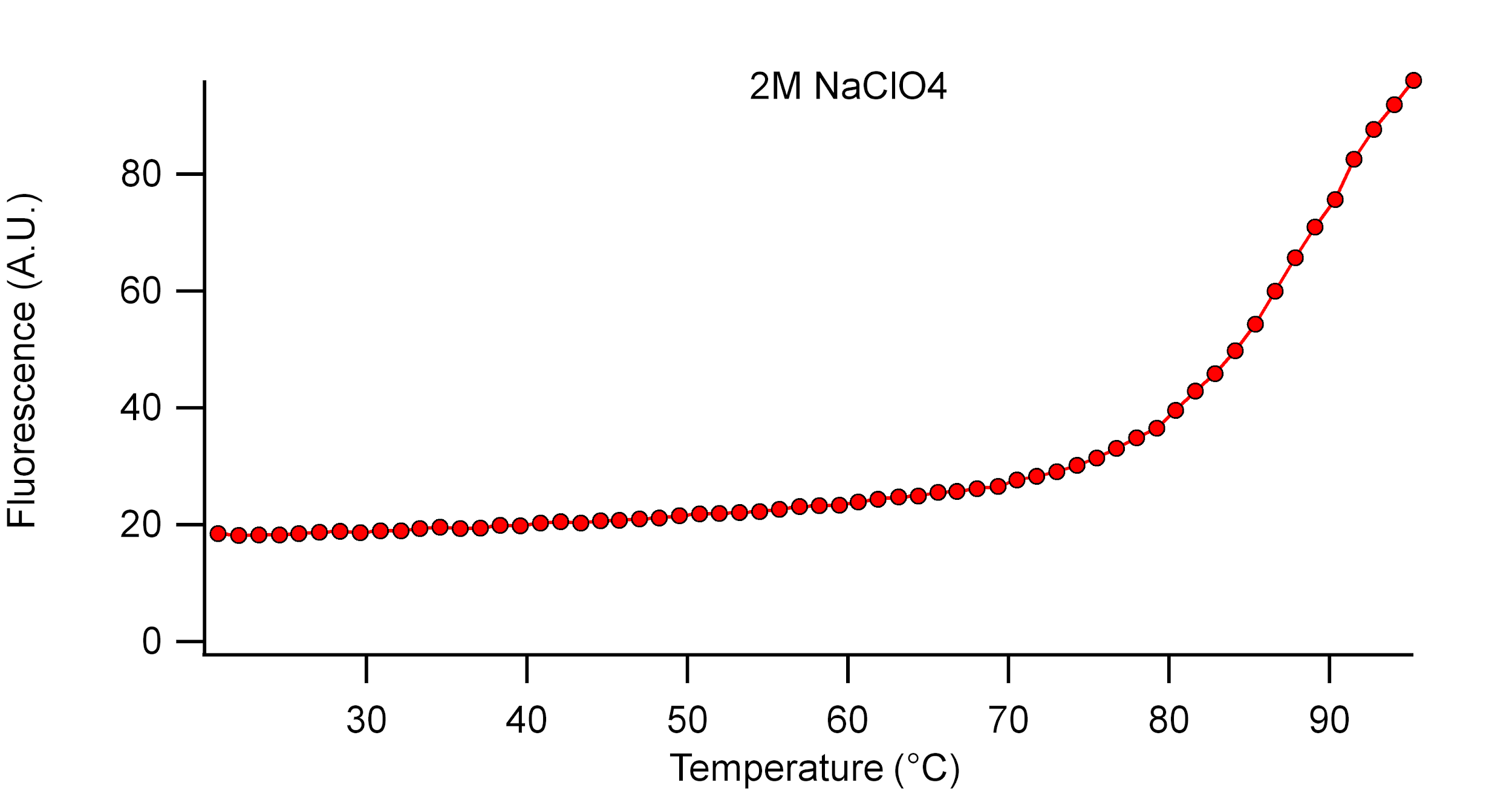

Supplementary Figure 11. Fluorescence melting curve (second heating trace) of **Fluorescein-G4-Quencher** in 3 M NaClO_4_.

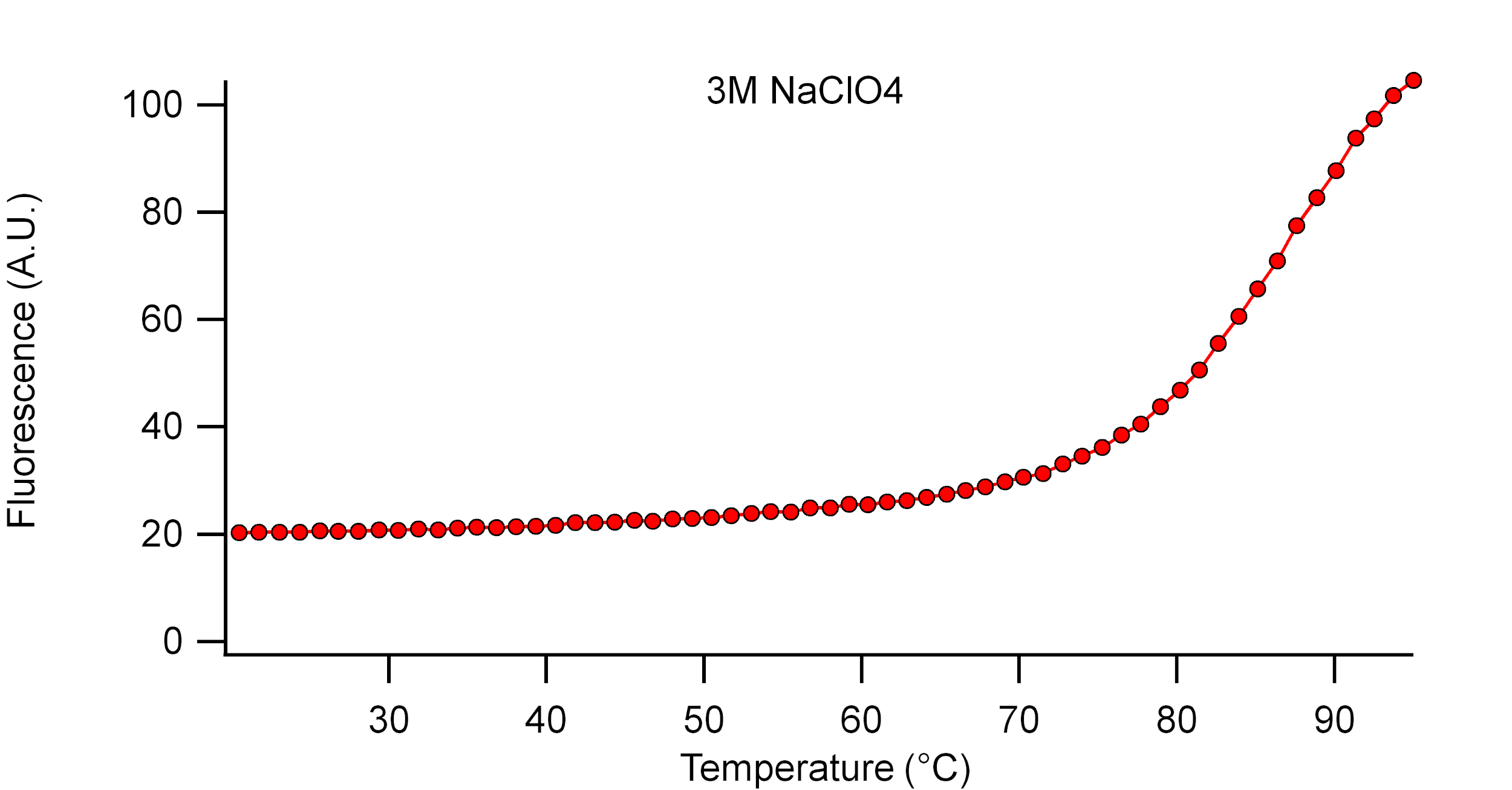

Supplementary Figure 12. Fluorescence melting curve (second heating trace) of **Fluorescein-G4-Quencher** in 4 M NaClO_4_.

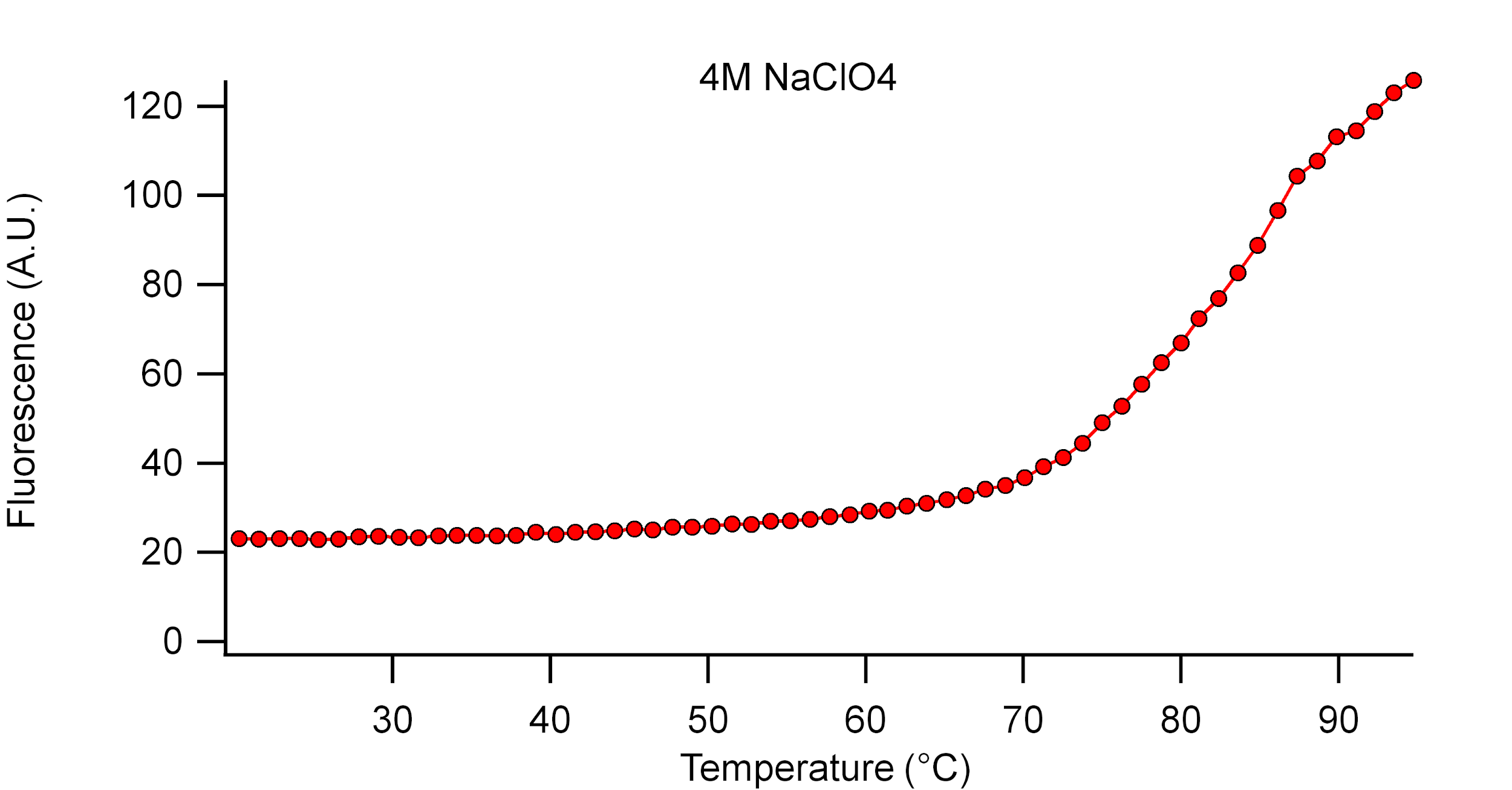

Supplementary Figure 13. Fluorescence melting curve (second heating trace) of **Fluorescein-G4-Quencher** in 5 M NaClO_4_.

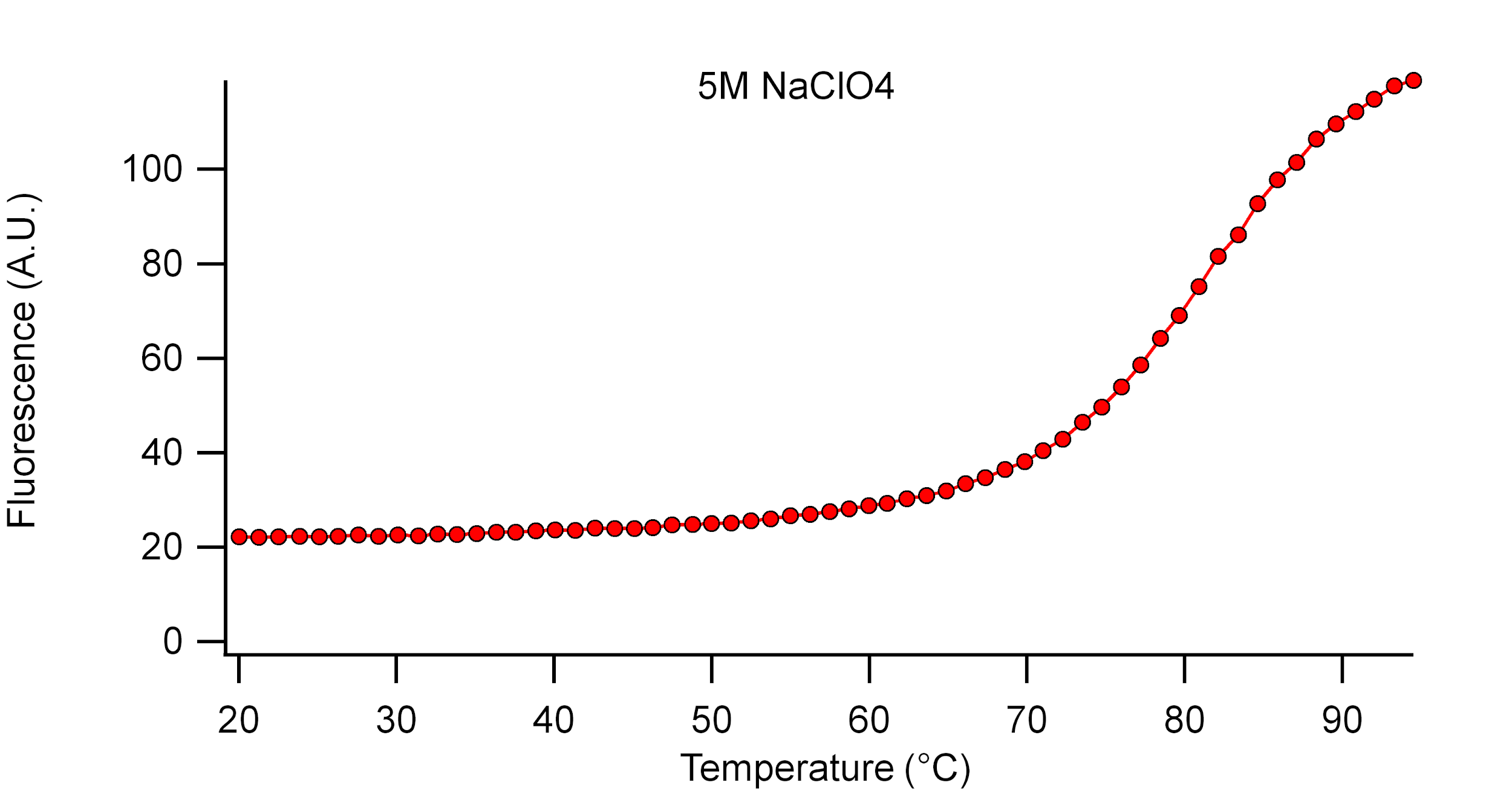

Supplementary Figure 14. Fluorescence melting curve (second heating trace) of **Fluorescein-G4-Quencher** in 6 M NaClO_4_.

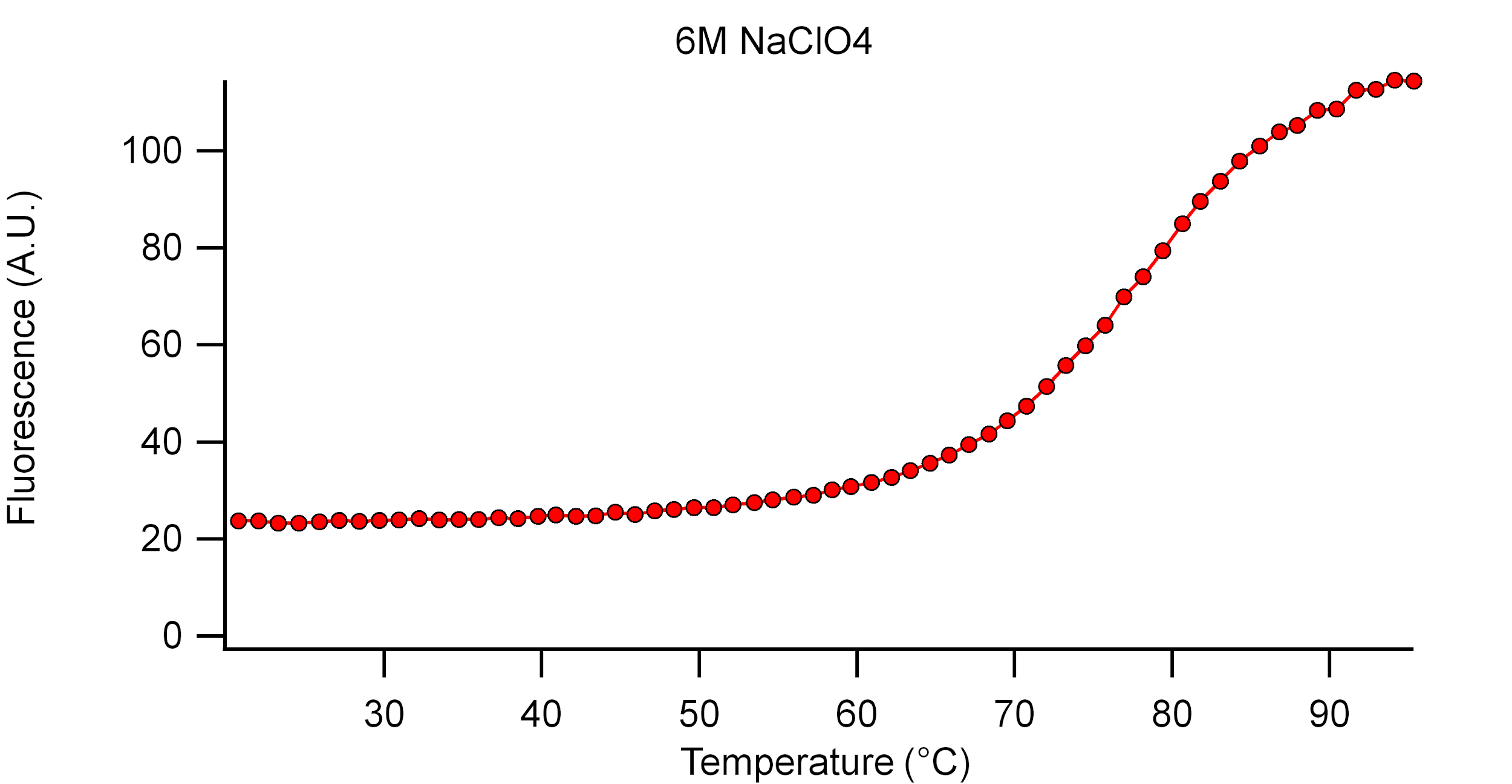

Supplementary Figure 15. Fluorescence melting curve (second heating trace) of **Fluorescein-G4-Quencher** in 7 M NaClO_4_.

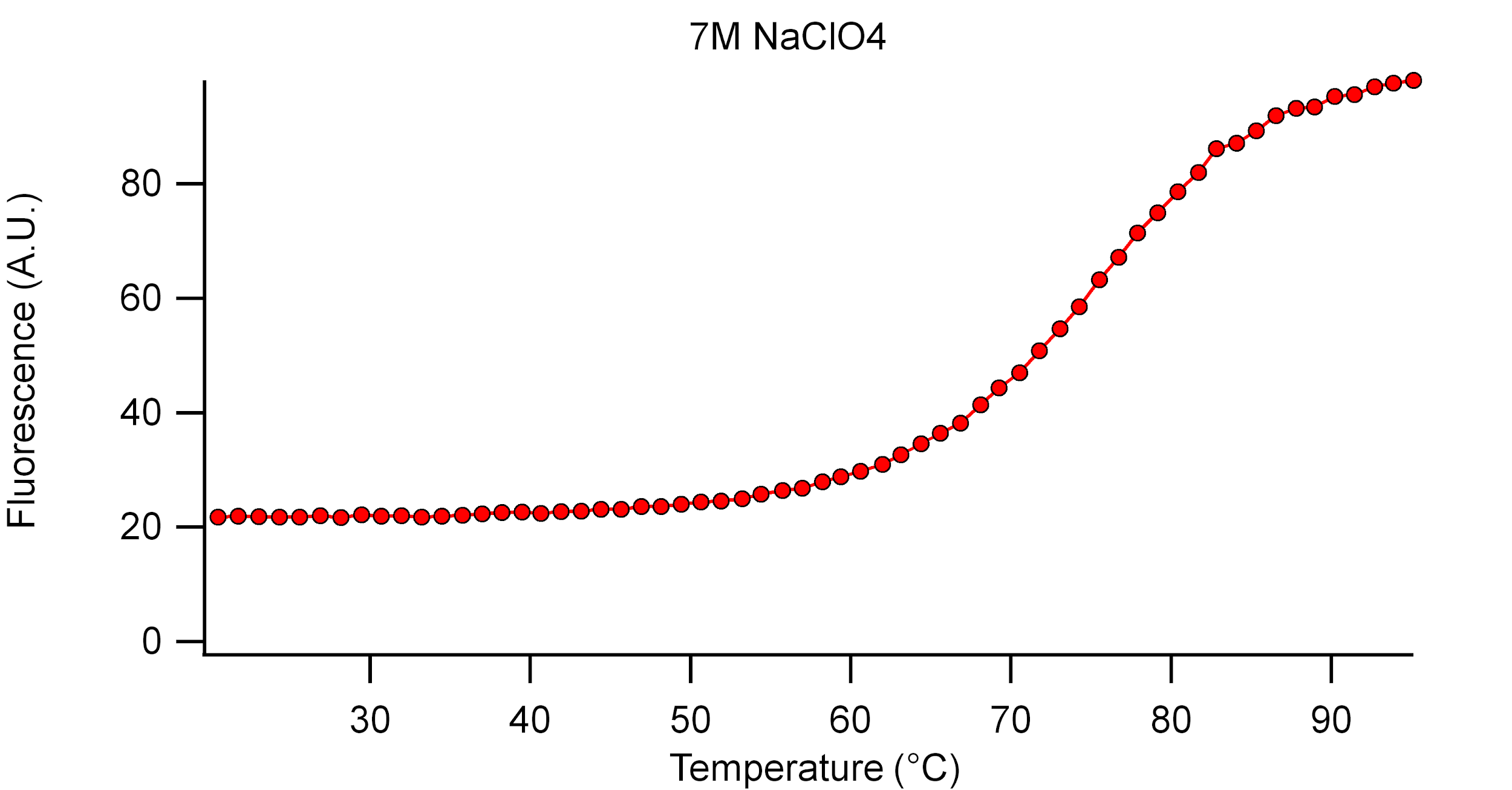

Supplementary Figure 16. Fluorescence melting curve (second heating trace) of **Fluorescein-G4-Quencher** in 8 M NaClO_4_.

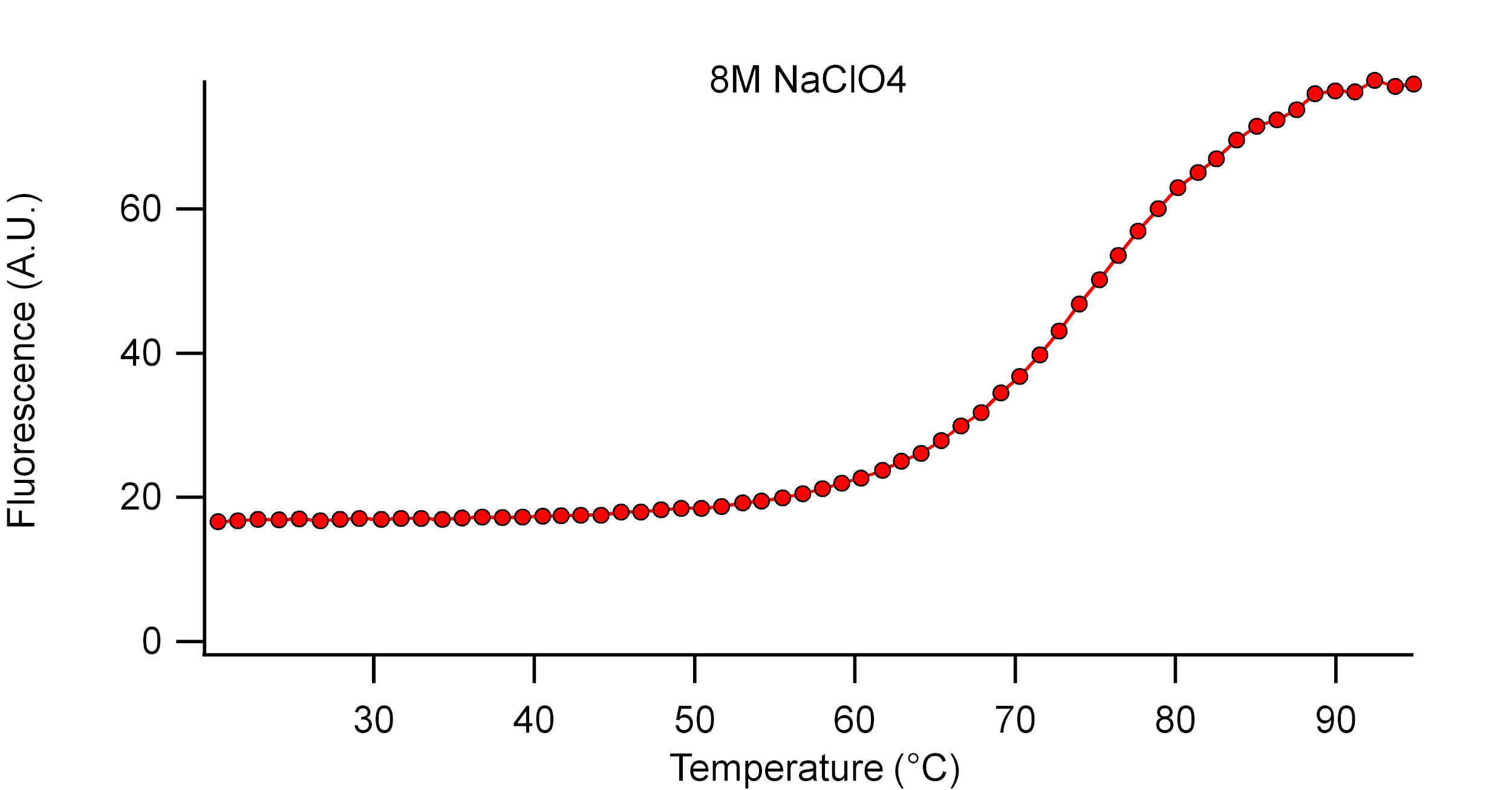

Supplementary Figure 17. Fluorescence melting curve (second heating trace) of **Fluorescein-G4-Quencher** in Saturated (ca. 9 M) NaClO_4_.

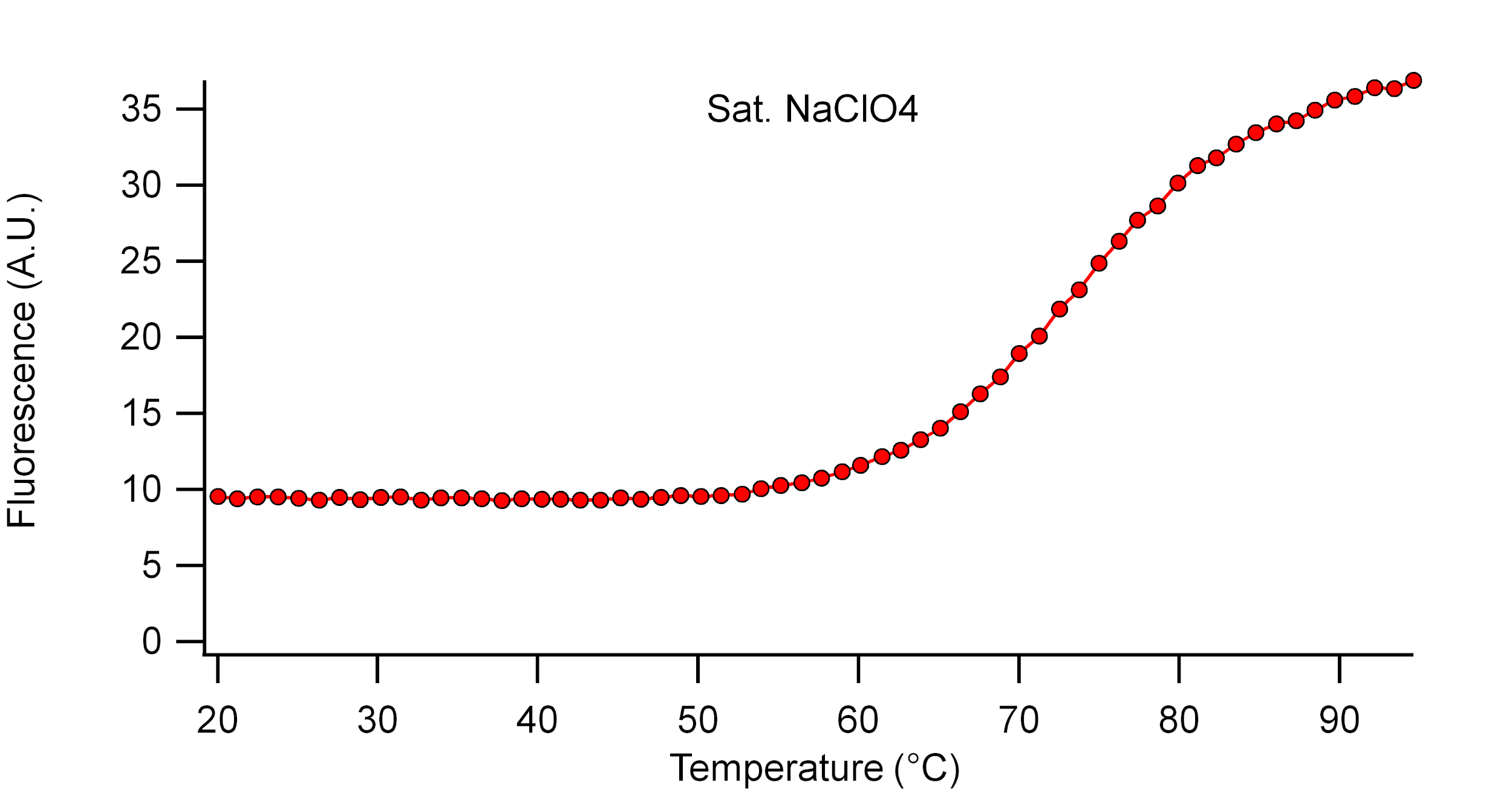

Supplementary Figure 18. Fluorescence melting curve (second heating trace) of **G4-Dark** in 0.1 M NaClO_4_.

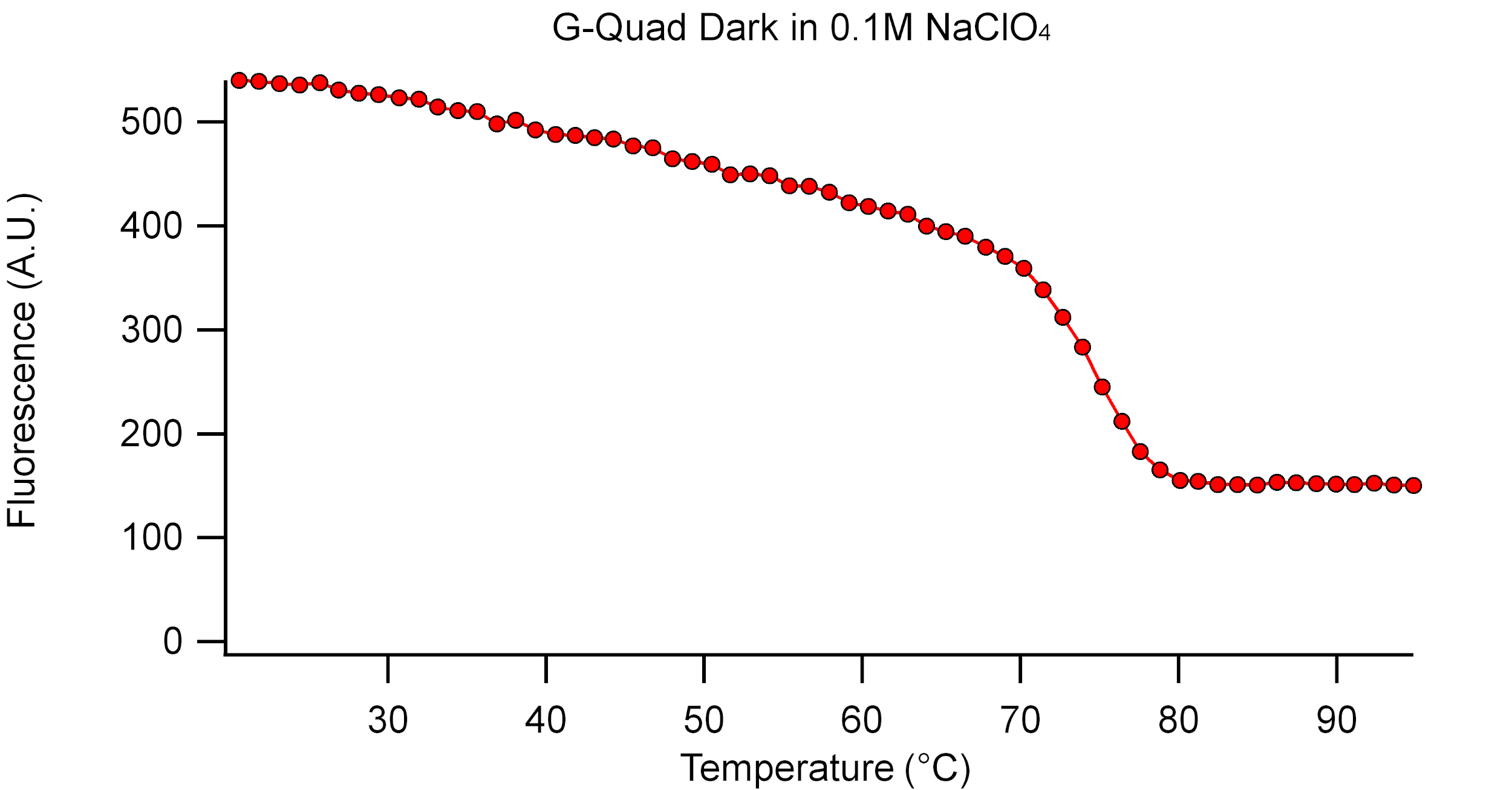

Supplementary Figure 19. Fluorescence melting curve (second heating trace) of **G4-Dark** in 0.5 M NaClO_4_.

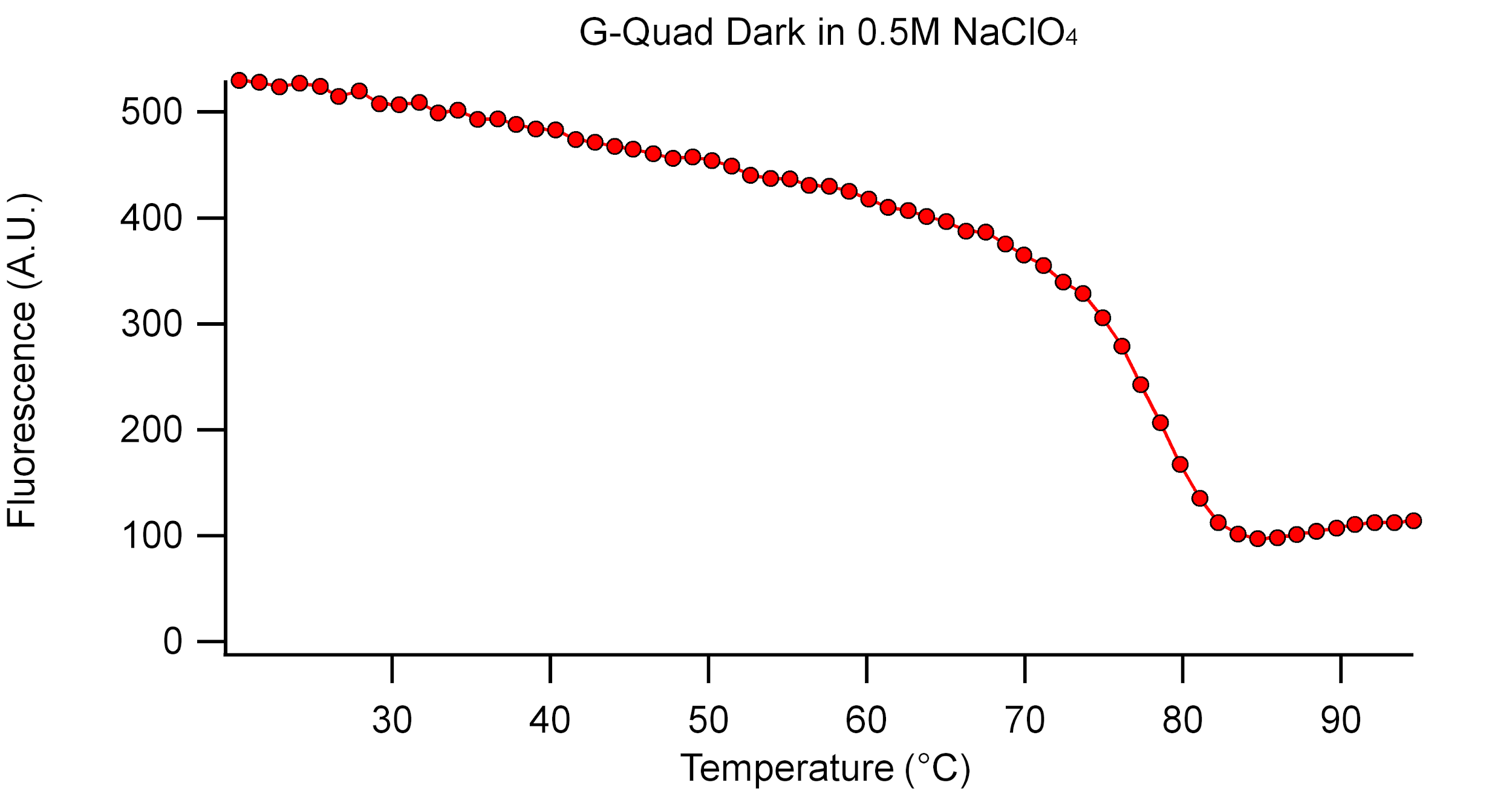

Supplementary Figure 20. Fluorescence melting curve (second heating trace) of **G4-Dark** in 1 M NaClO_4_.

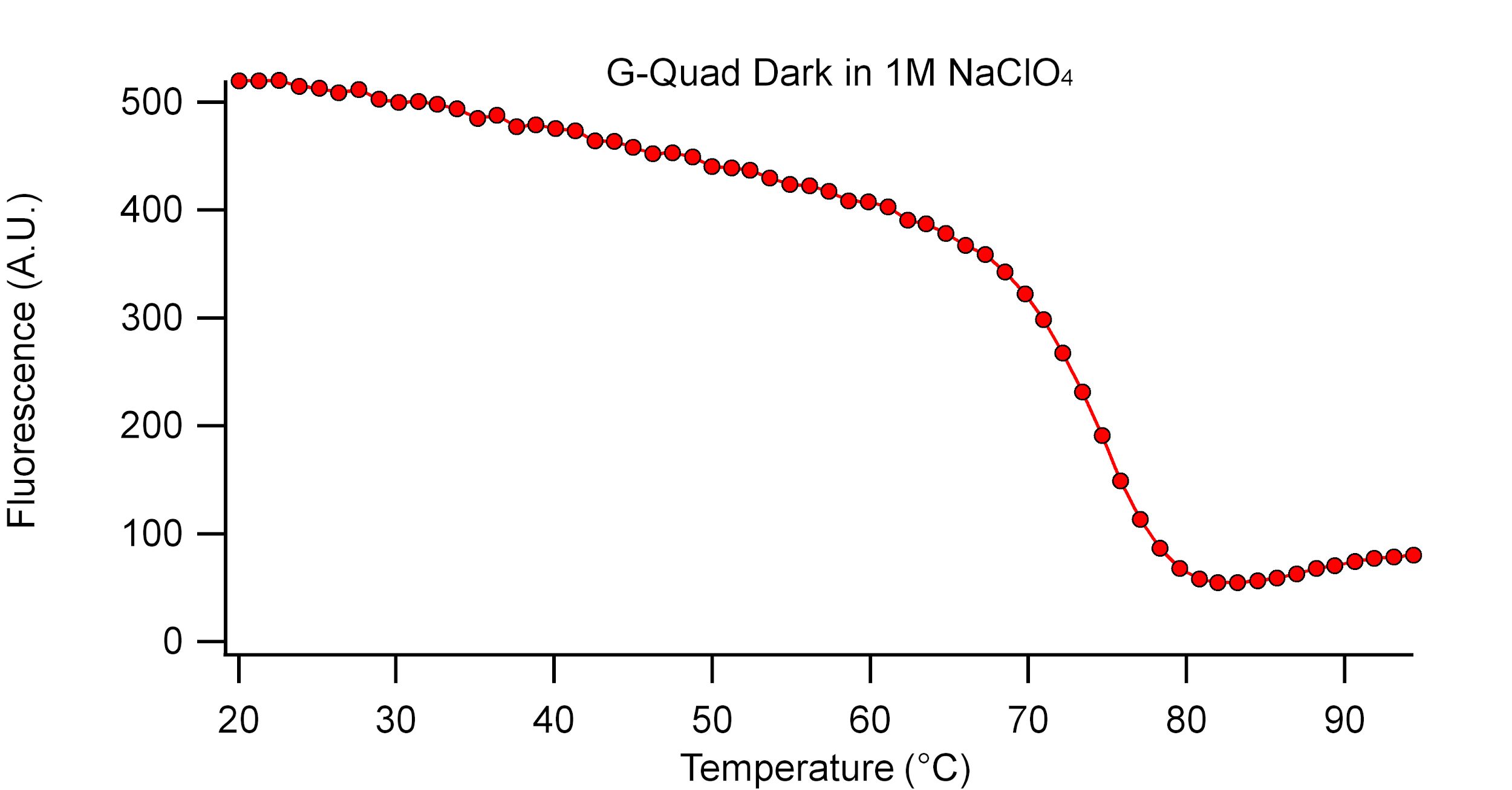

Supplementary Figure 21. Fluorescence melting curve (second heating trace) of **G4-Dark** in 2 M NaClO_4_.

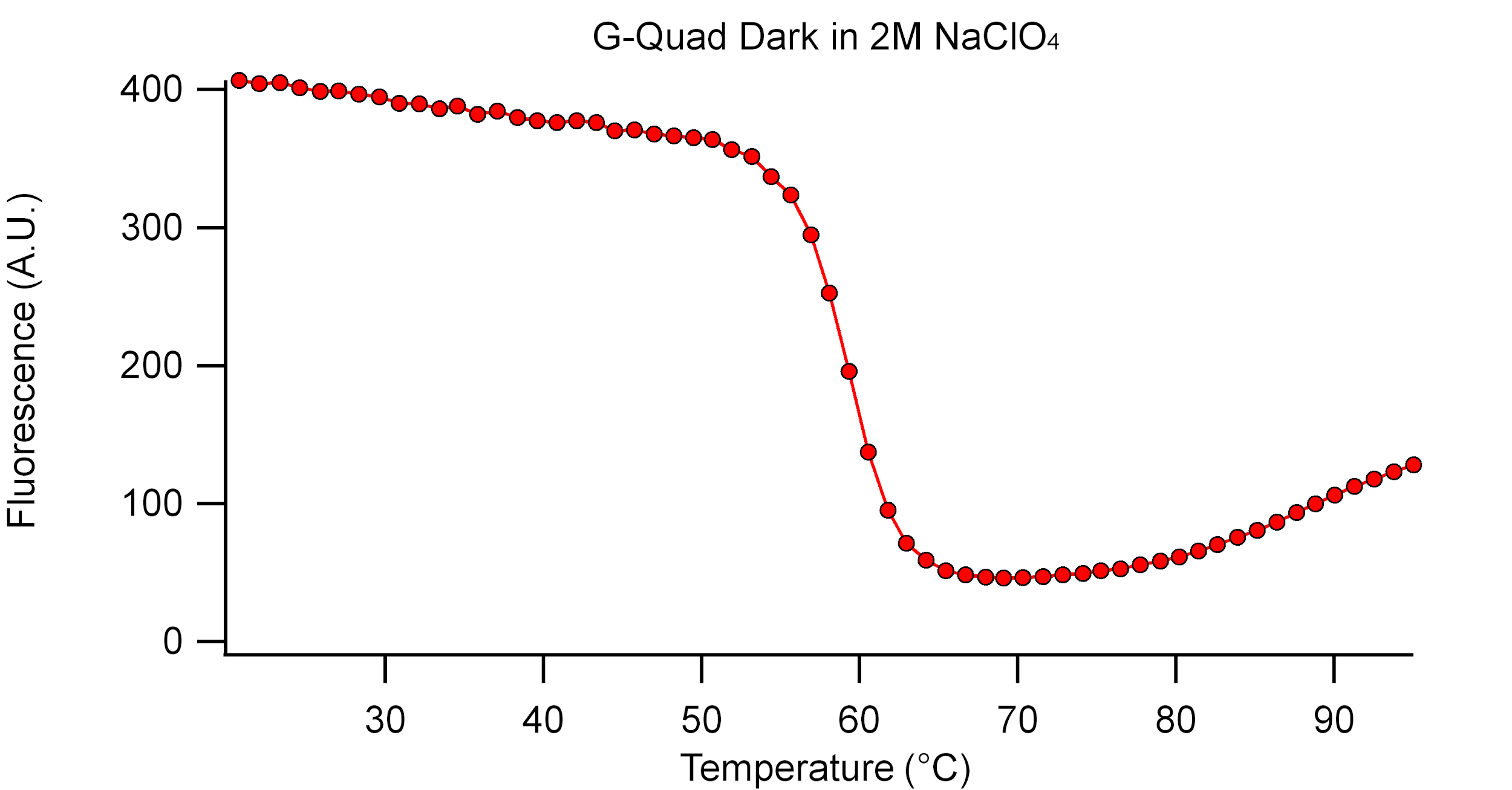

Supplementary Figure 22. Fluorescence melting curve (second heating trace) of **G4-Dark** in 3 M NaClO_4_.

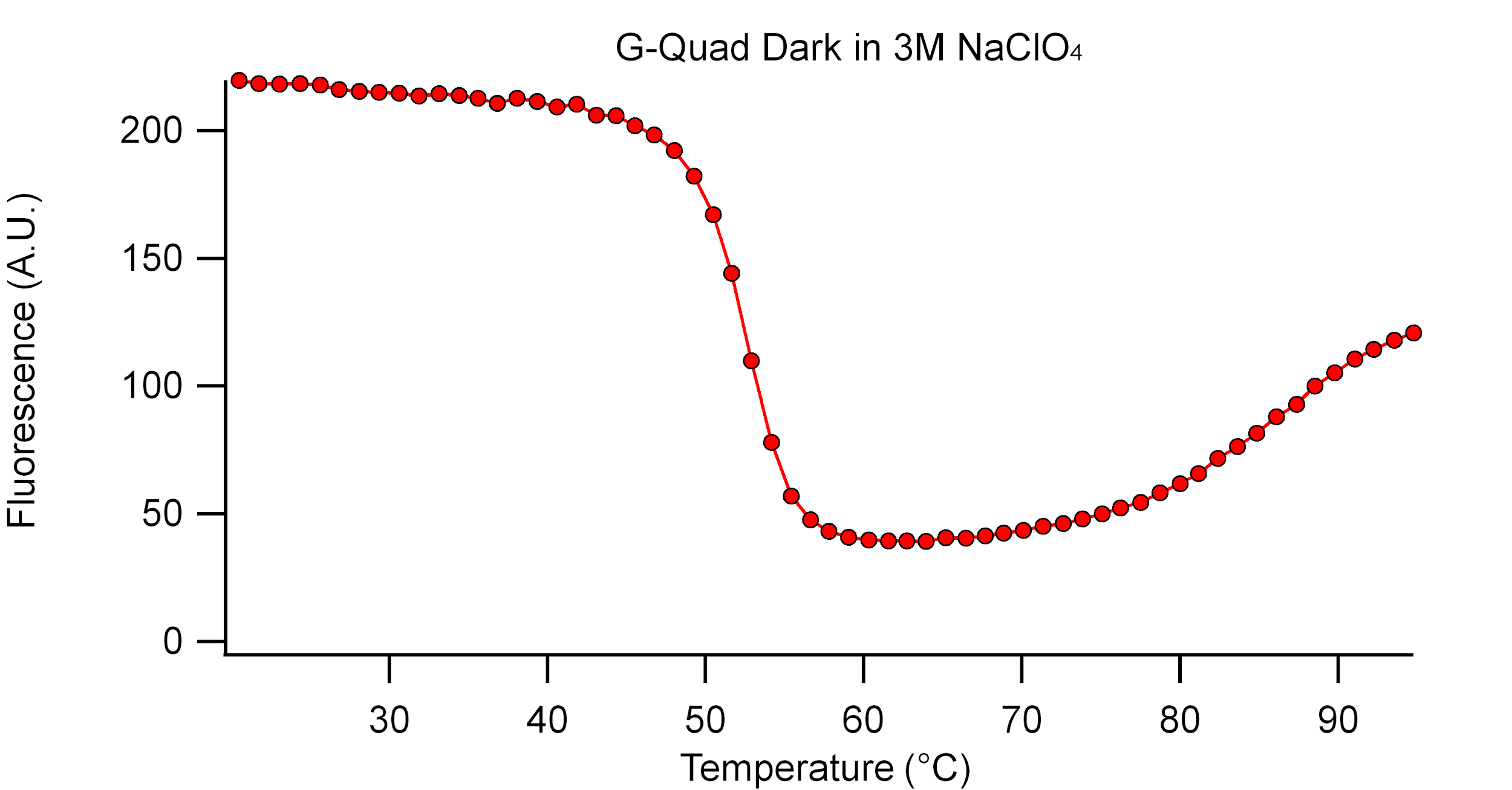

Supplementary Figure 23. Fluorescence melting curve (second heating trace) of **G4-Dark** in 4 M NaClO_4_.

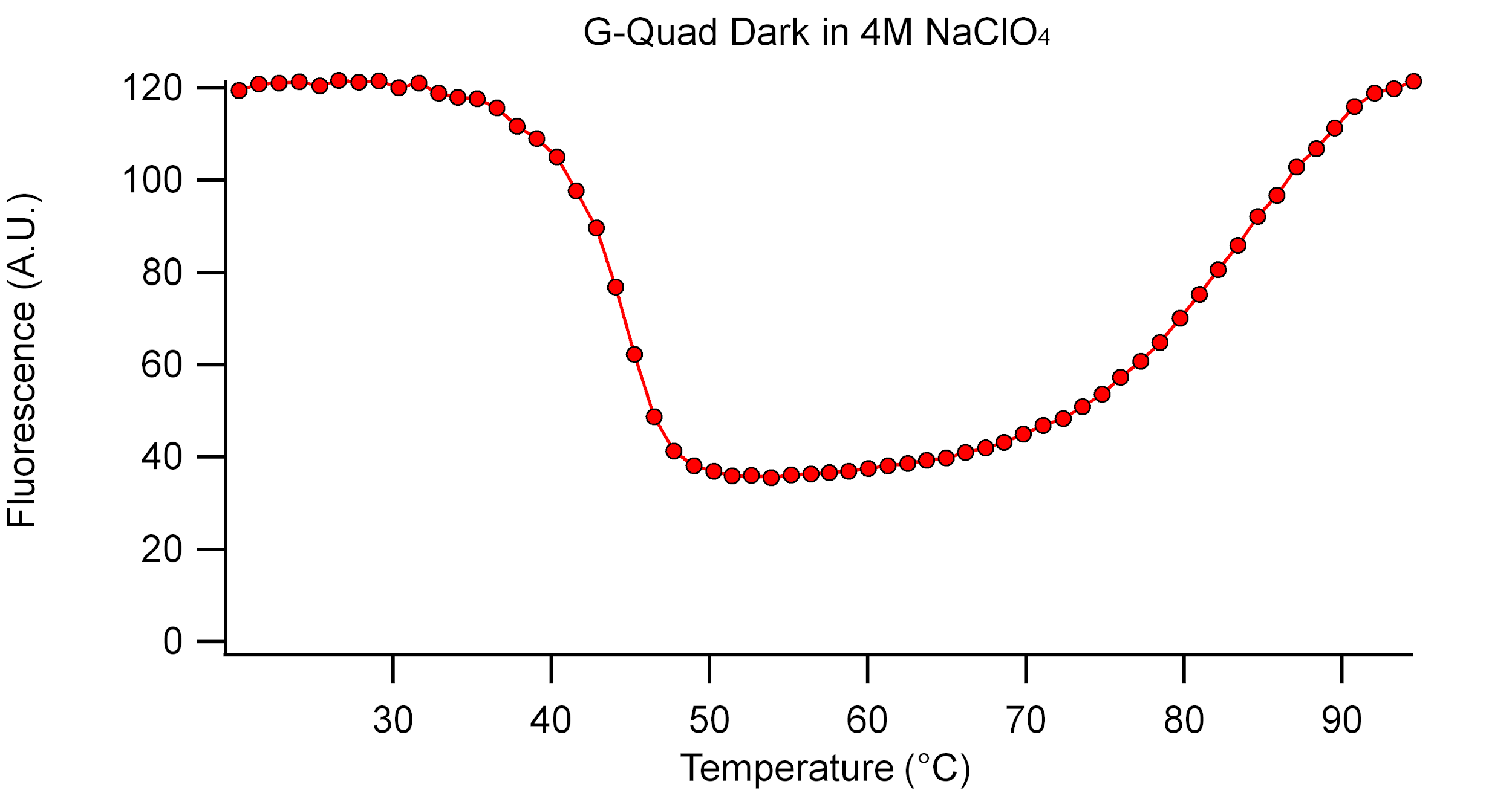

Supplementary Figure 24. Fluorescence melting curve (second heating trace) of **G4-Dark** in 5 M NaClO_4_.

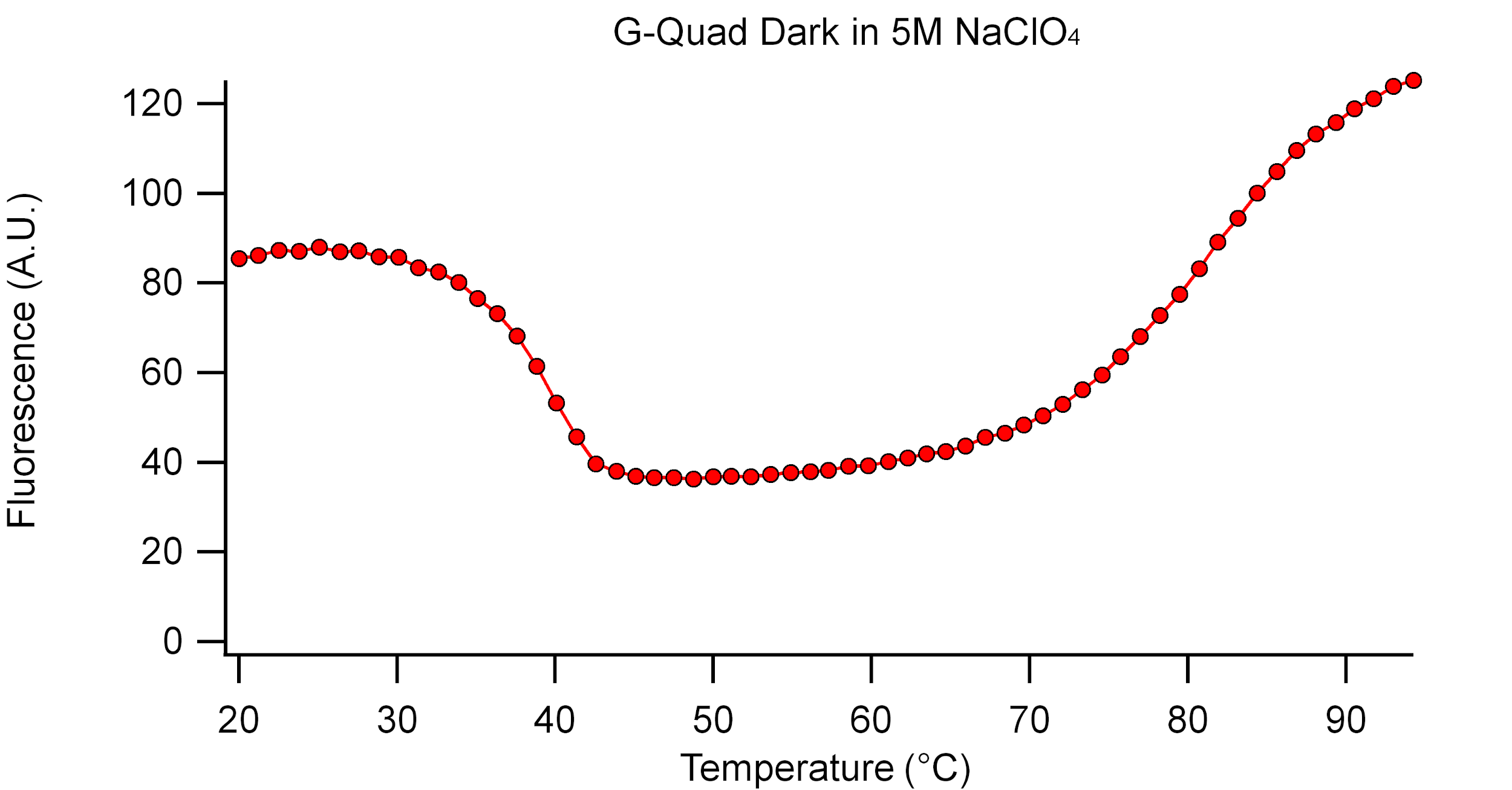

Supplementary Figure 25. Fluorescence melting curve (second heating trace) of **G4-Dark** in 6 M NaClO_4_.

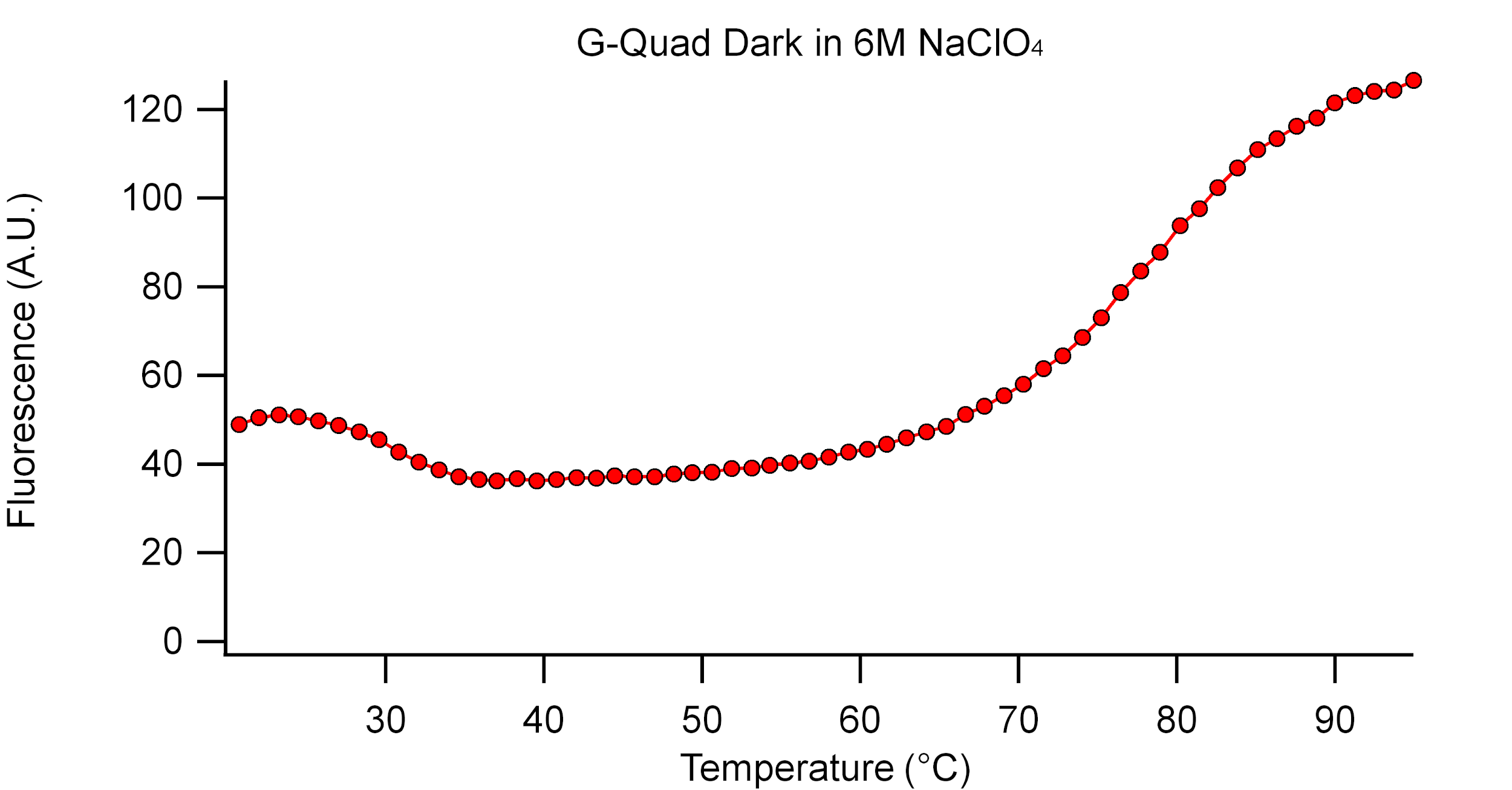

Supplementary Figure 26. Fluorescence melting curve (second heating trace) of **G4-Dark** in 7 M NaClO_4_.

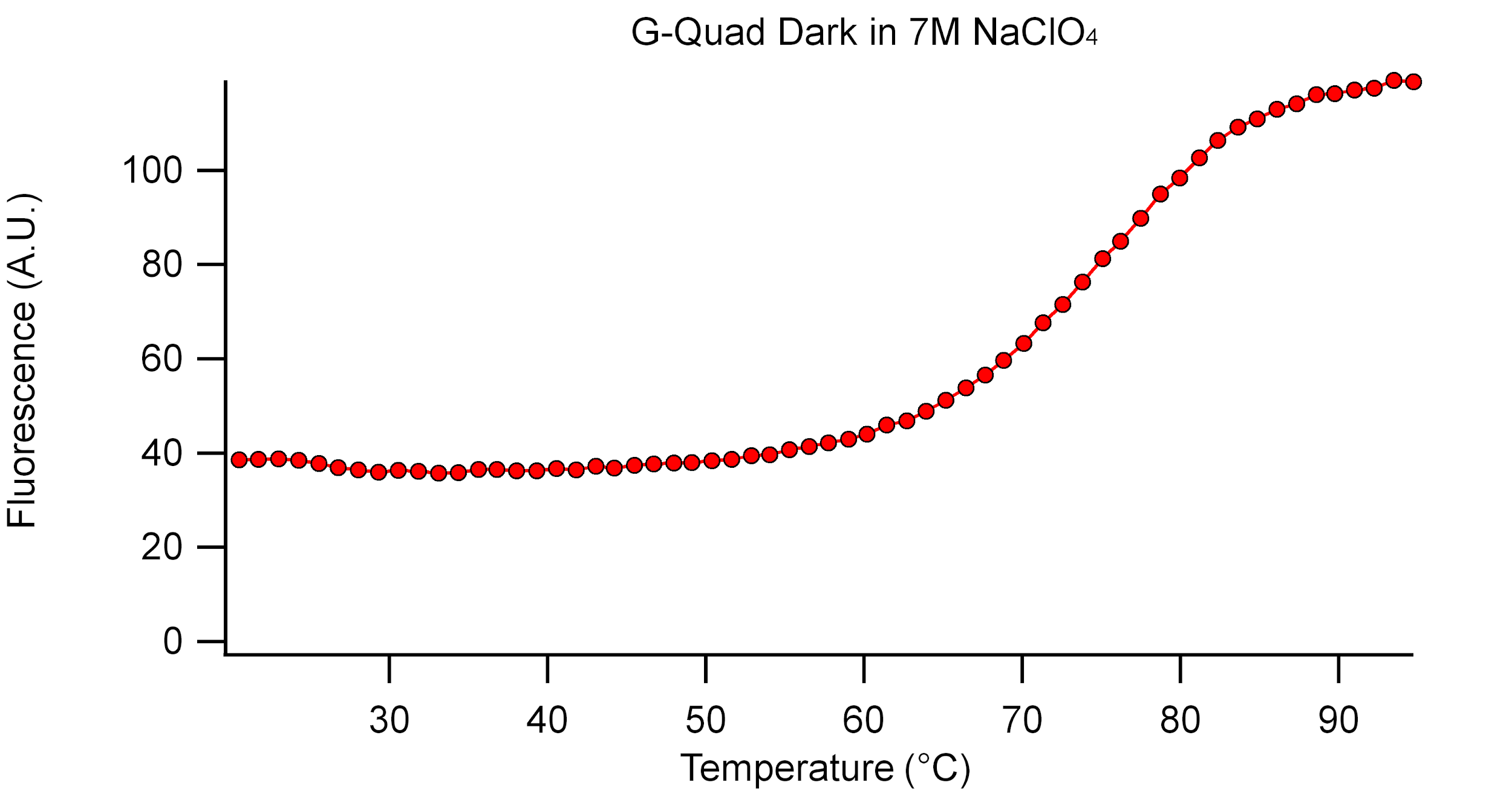

Supplementary Figure 27. Fluorescence melting curve (second heating trace) of **G4-Dark** in 8 M NaClO_4_.

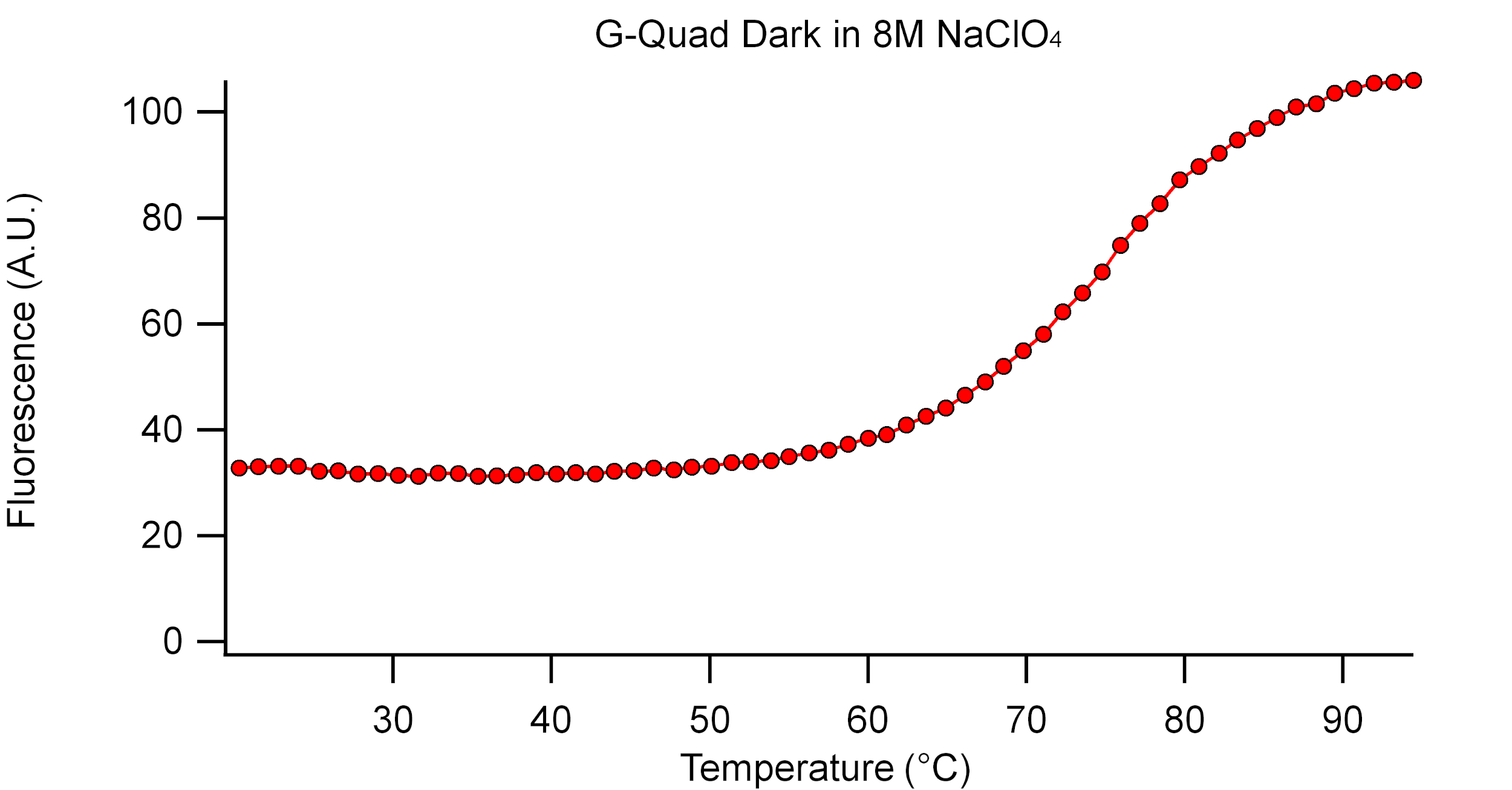

Supplementary Figure 28. Fluorescence melting curve (second heating trace) of **G4-Dark** in Saturated NaClO_4_.

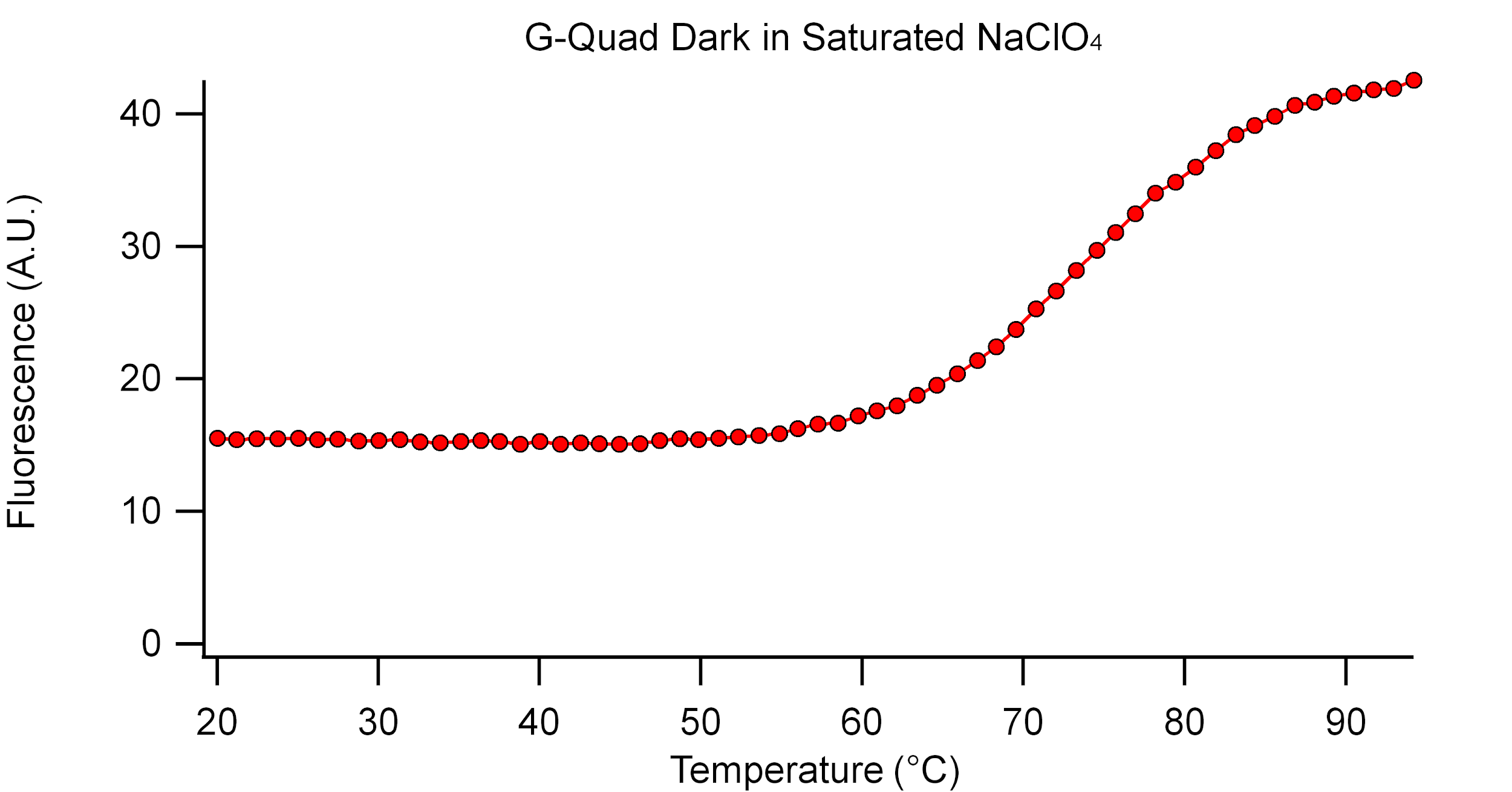

Supplementary Figure 29. Engineering drawing of 3D printed tube holder/camera mount component of imaging jig. Dimensions in mm.

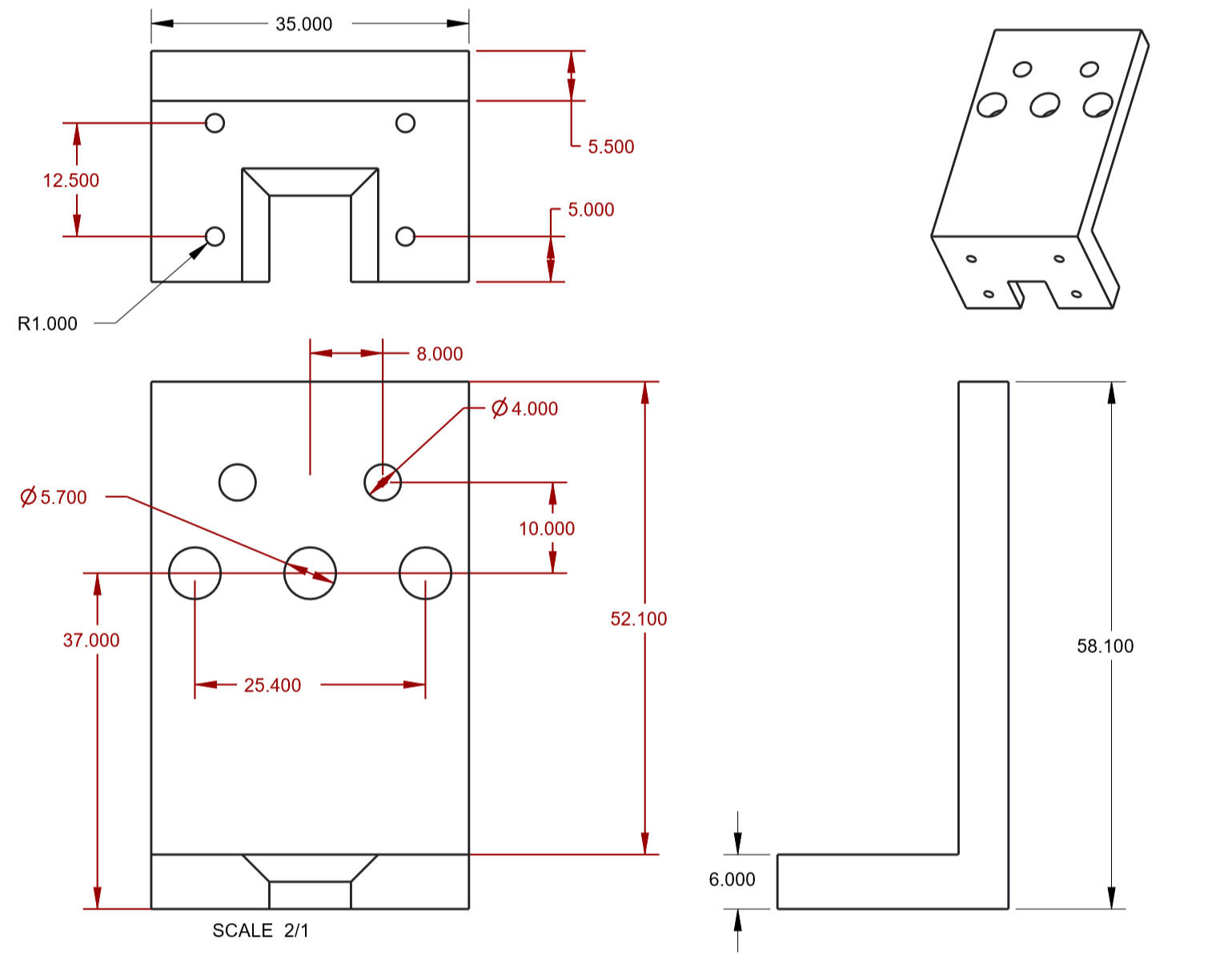

Supplementary Figure 30. Engineering drawing of 3D printed led holder component of imaging jig. Dimensions in mm.
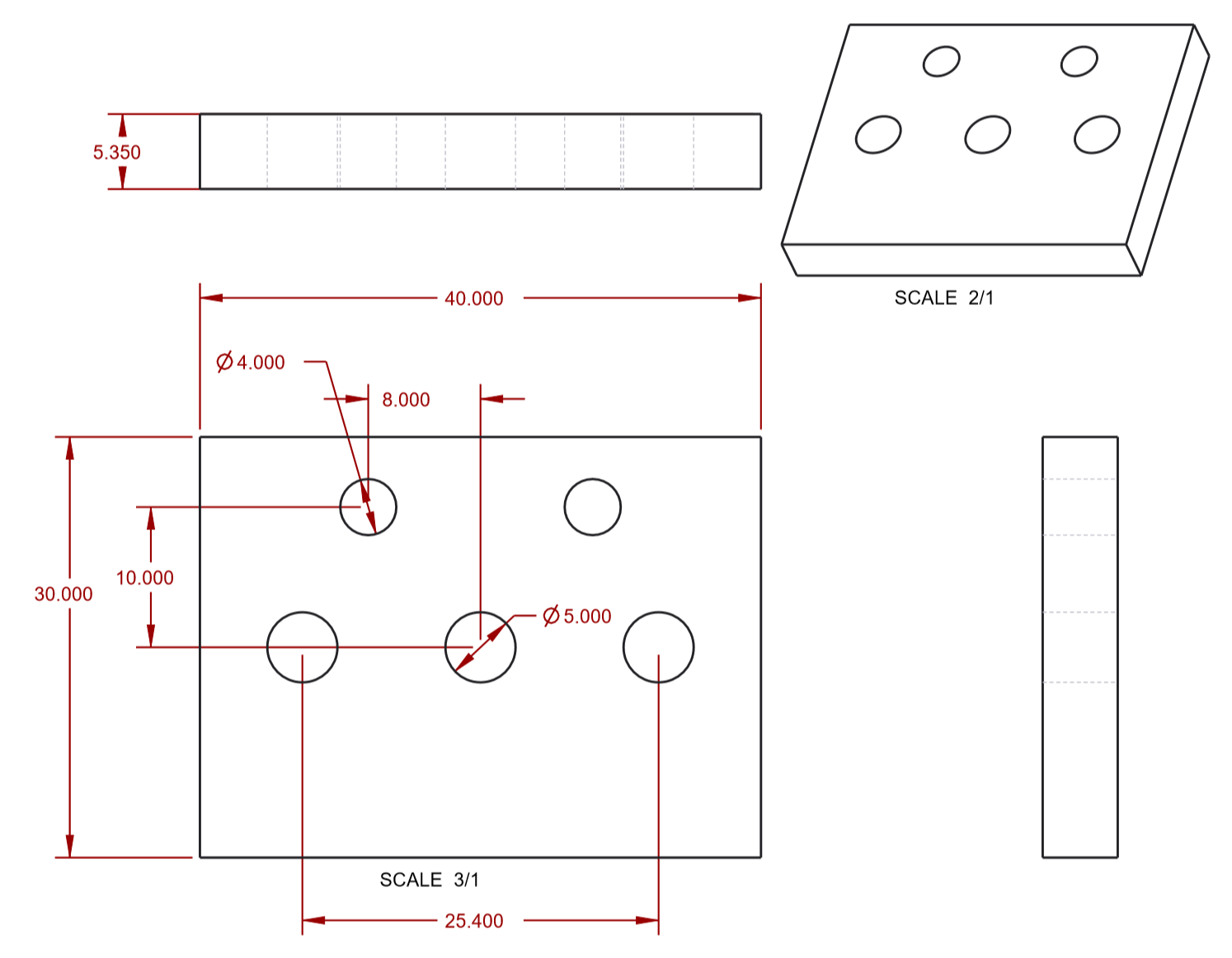

Supplementary Figure 31. CAD Mockup of Fluorescence Imaging Jig

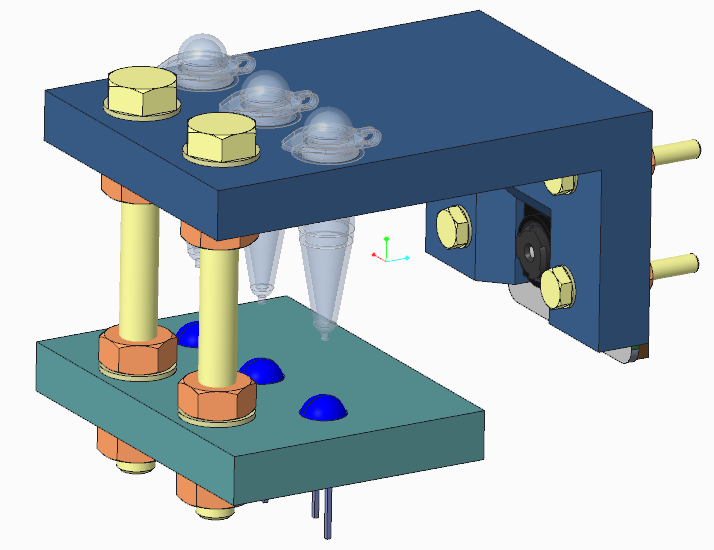

Supplementary Figure 32. CAD Mockup of Fluorescence Imaging Jig (Alternate view)

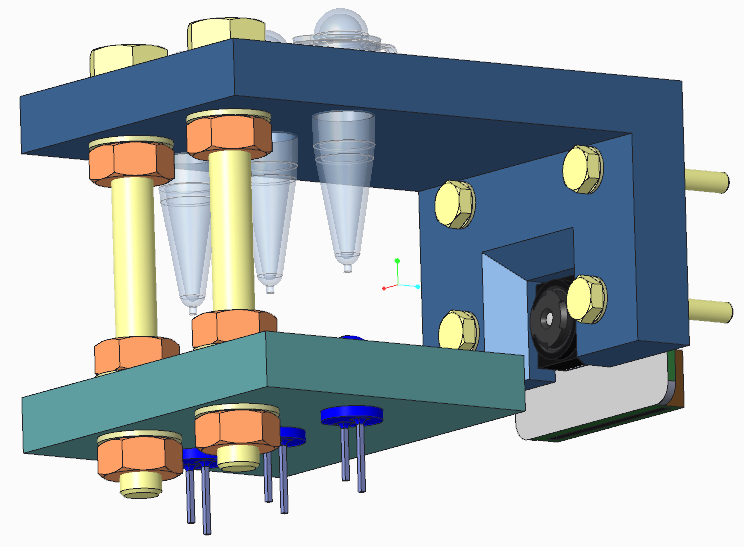

Supplementary Figure 33. Imaging jig with tubes.

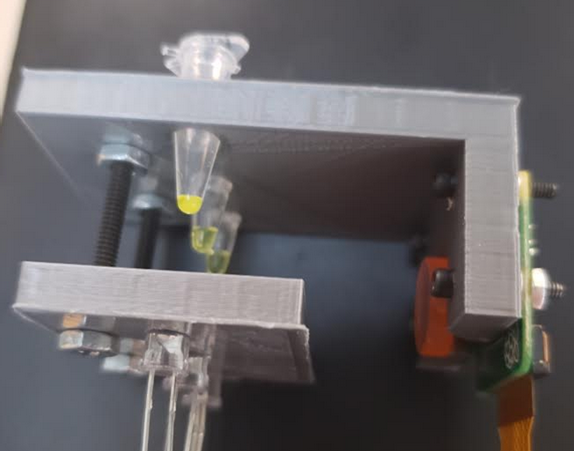

Supplementary Figure 34. Imaging jig (top view).

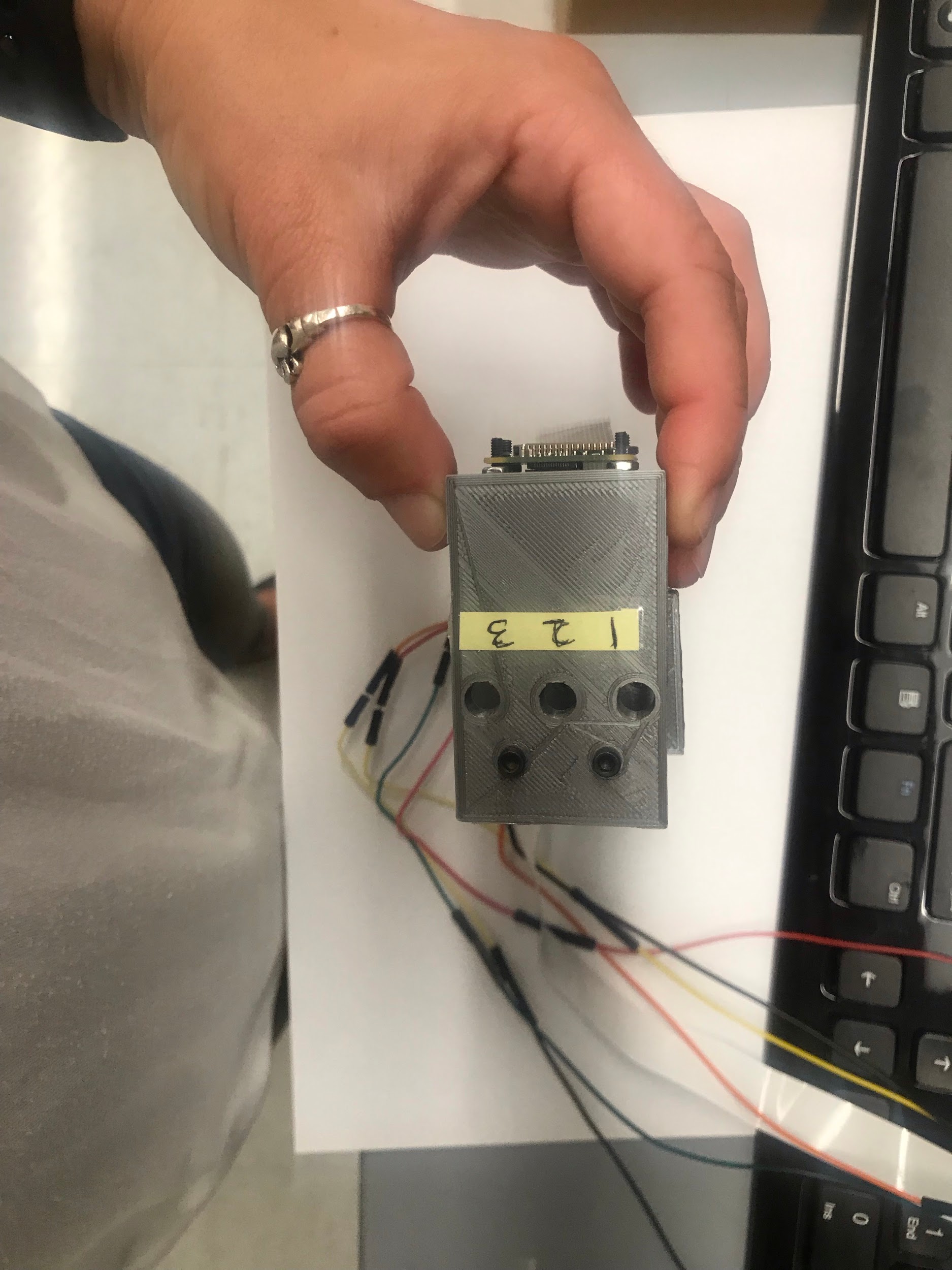

Supplementary Figure 35. Imaging Jig (side view)
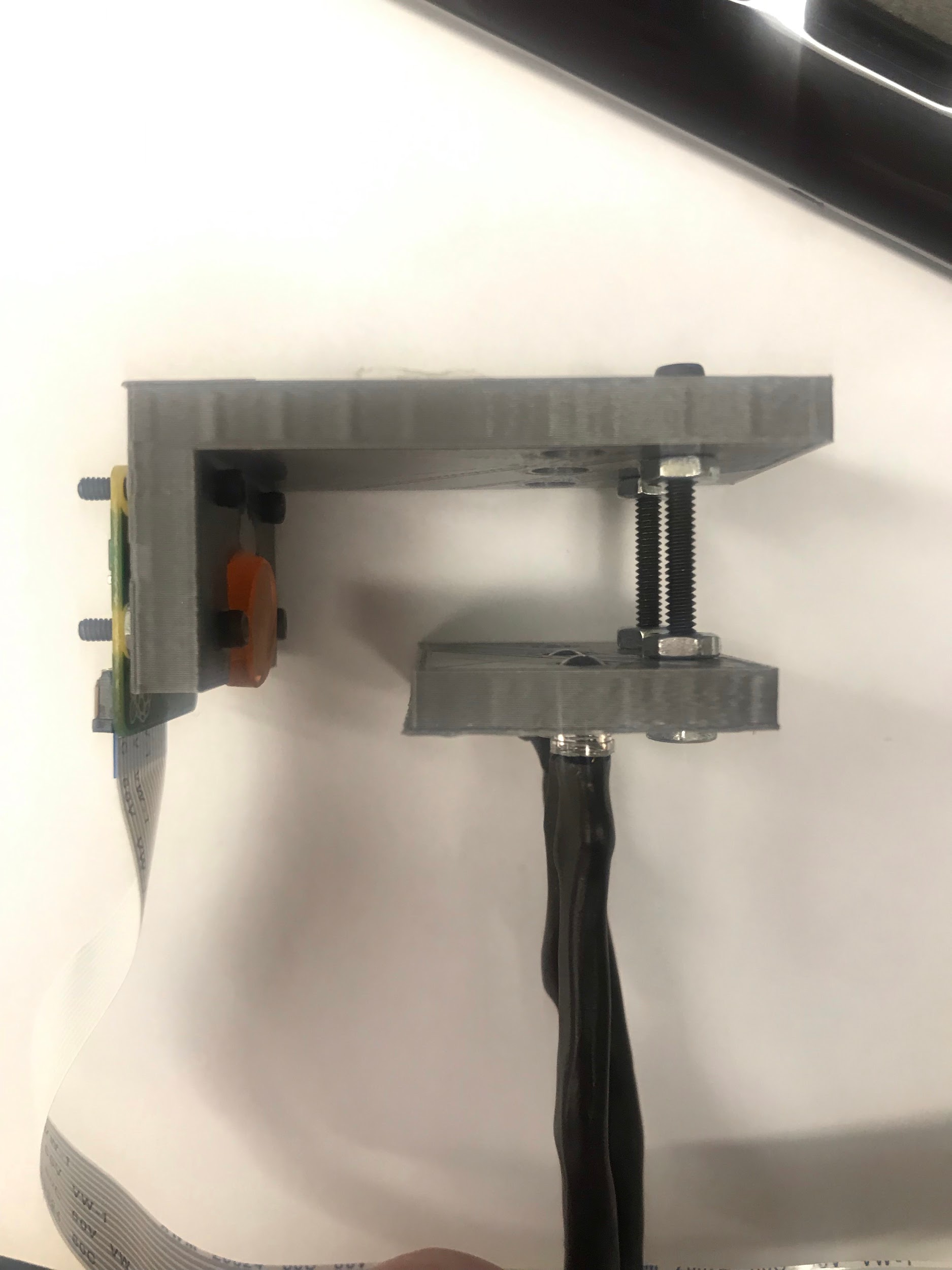

Supplementary Figure 36. Imaging Jig (front view).

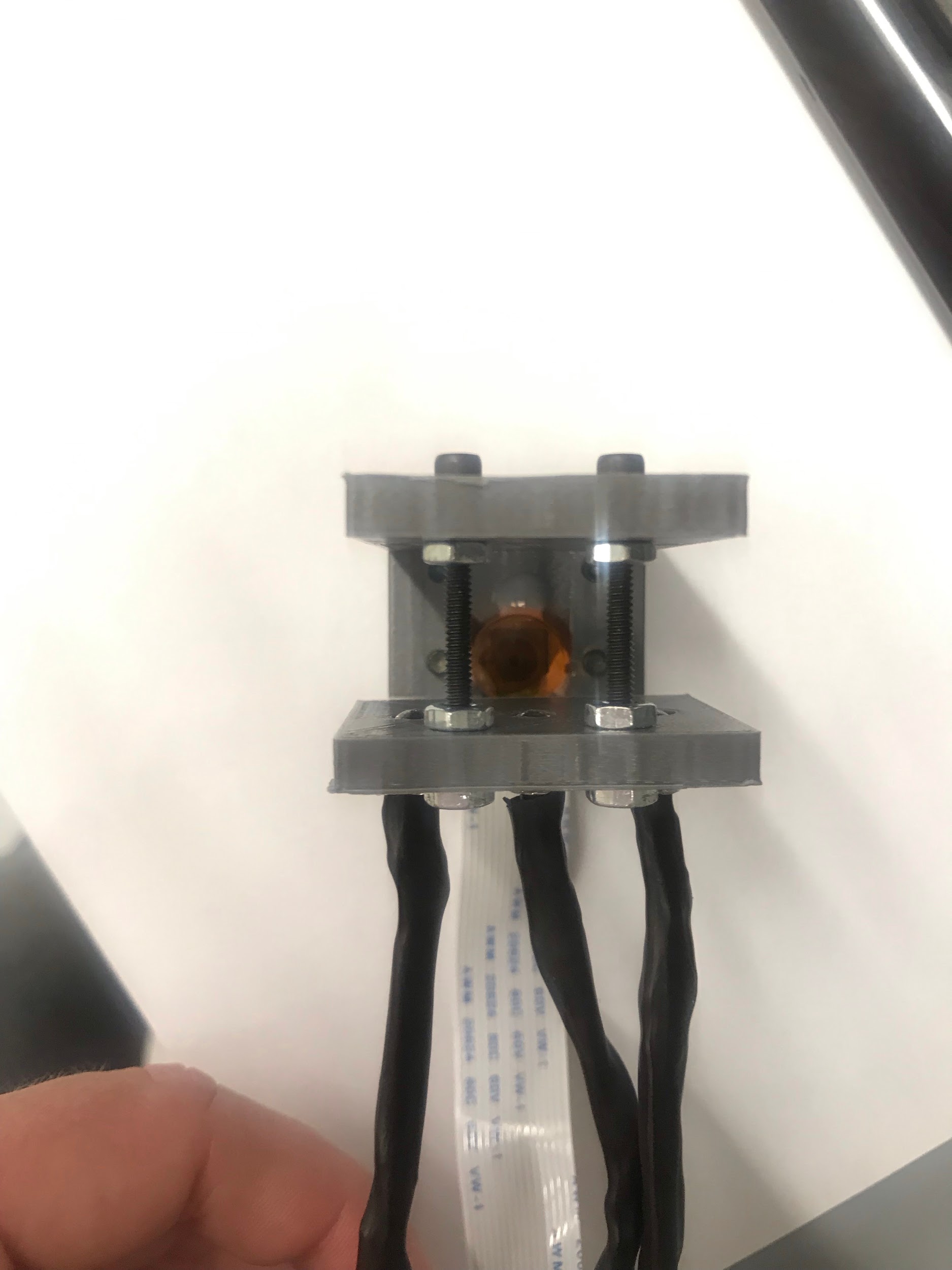

Supplementary Video 1. Fluorescence Imaging shows G4-Dark and Duplex-Dark switching structure under vacuum control with real-time plotting of fluorescence intensity. Available online.

Supplementary Video 2. Fluorescence Imaging shows G4-Dark and Duplex-Dark switching structure under vacuum control (With annotations and captions). Available online.

Supplementary Code 1. Python code for controlling Raspberry Pi for reaction monitoring and overlaying plots of fluorescence vs. time.

#!/usr/bin/env python3

\# -*- coding: utf-8 -*-

"""

Created on Wed Jun 20 11:16:14 2018

@author: Lauren Aufdembrink

"""

#####################################Code starts below here#########################################

#@author: Lauren Aufdembrink

import time

from picamera import PiCamera

from gpiozero import LED

import numpy as np

from PIL import Image

import matplotlib.pyplot as plt

import os

Minutes=60*float(input('How long you would like to run in hours:'))

Intervals=float(input('How often do you want it to take photos in minutes:'))

PicturesTake=int(Minutes/Intervals)

Sample1=input('Name of sample in Slot 1:')

Sample2=input('Name of sample in Slot 2:')

Sample3=input('Name of sample in Slot 3:')

Data=input('Name file to save pixel intensities:')

ImageFolder=input('Name a folder to save tube images:')

PlotFolder=input('Name a folder to save plot images:')

OverlayFolder=input('Name a folder to save overlay images:')

pause=input('Press enter when you are ready to begin')

os.makedirs(ImageFolder)

os.makedirs(PlotFolder)

os.makedirs(OverlayFolder)

#######My function############

def PixelAvg(image, Slot1, Slot2, Slot3):

#loading the image in:

im=Image.open(image)

#imarray=np.array(im)

#to take a subset from image for each tube

#Slot1=imarray[50:240,1:427]

#Slot2=imarray[52:240,427:854]

#Slot3=imarray[50:240,855:1279]

#to go from an array back to an image:

#this is only necessary if you need to view your subset:

#Slot1Im=Image.fromarray(Slot1)

#Slot2Im=Image.fromarray(Slot2)

#Slot3Im=Image.fromarray(Slot3)

#converting Images to grayscale:

Slot1Gray=np.array(im.convert('L'),'f')[50:240,1:427]

Slot2Gray=np.array(im.convert('L'),'f')[52:240,427:854]

Slot3Gray=np.array(im.convert('L'),'f')[50:240,855:1279]

#finding the index of Max value of the image

#this will use that bright dot at the bottom of the tube to base my pixel subset off of

Slot1MaxInd=np.unravel_index(np.argmax(Slot1Gray), (190,426))

Slot2MaxInd=np.unravel_index(np.argmax(Slot2Gray), (188,427))

Slot3MaxInd=np.unravel_index(np.argmax(Slot3Gray), (190,424))

#after finding the max value indexes above, taking the subset to average over:

Slot1Pixel=Slot1Gray[(Slot1MaxInd[0]-30):(Slot1MaxInd[0]-10),(Slot1MaxInd[1]-10):(Slot1MaxInd[1]+10)]

Slot2Pixel=Slot2Gray[(Slot2MaxInd[0]-30):(Slot2MaxInd[0]-10),(Slot2MaxInd[1]-10):(Slot2MaxInd[1]+10)]

Slot3Pixel=Slot3Gray[(Slot3MaxInd[0]-30):(Slot3MaxInd[0]-10),(Slot3MaxInd[1]-10):(Slot3MaxInd[1]+10)]

#now find the mean of your pixel selection:

S1mean=np.mean(Slot1Pixel)

S2mean=np.mean(Slot2Pixel)

S3mean=np.mean(Slot3Pixel)

Slot1.append(S1mean)

Slot2.append(S2mean)

Slot3.append(S3mean)

return(S1mean, S2mean, S3mean, Slot1, Slot2, Slot3);

def ImageView(TubePic,PlotPic,i):

background= Image.open(TubePic)

overlay= Image.open(PlotPic)

background.paste(overlay, (400,360))

background.save(OverlayFolder+'/Ovly%s.jpg'%i, 'jpeg')

return(background);

#insert the name of the file where the data is being stored once analyzed

DataFile=open(Data,'a+')

led= LED(12)

led2= LED(13)

led3= LED(19)

led.on()

led2.on()

led3.on()

camera=PiCamera()

camera.resolution=(1280,720)

camera.framerate=1

time.sleep(2)

camera.shutter_speed=750000

camera.exposure_mode='off'

camera.awb_mode='off'

camera.awb_gains=[3,1] #{red,blue]

camera.iso=100

time.sleep(2)

finalline='0 0 0'

Slot1=[]

Slot2=[]

Slot3=[]

Time=[]

led.off()

led2.off()

led3.off()

camera.stop_preview()

mydpi=100

for i in range(PicturesTake):

time.sleep((Intervals*60)-3.5)

Time.append(i*Intervals)

led.on()

led2.on()

led3.on()

camera.annotate_text=finalline

camera.capture(ImageFolder+'/Rxn%s.jpg'%i)

led.off()

led2.off()

led3.off()

values=PixelAvg(ImageFolder+'/Rxn%s.jpg'%i, Slot1, Slot2, Slot3)

S1=str(round(values[0]))

S2=str(round(values[1]))

S3=str(round(values[2]))

DataFile.write(S1+'\t'+S2+'\t'+S3+'\n')

finalline='S1:'+S1+'	S2:'+S2+'	S3:'+S3

plt.plot(Time, Slot1,'g-', label=Sample1)

plt.plot(Time, Slot2,'r--',label=Sample2)

plt.plot(Time, Slot3, 'k^:', label=Sample3)

plt.legend(loc=7, frameon=False)

plt.xlim(0,Time[i]+5)

plt.ylim(0,255)

plt.xlabel('Time')

plt.ylabel('Intensity (Au)')

plt.title('Fluorescence Monitoring in Vacuum Chamber')

plt.savefig(PlotFolder+'/Plot%s.jpg'%i, dpi=mydpi*0.75)

plt.close()

ImageView(ImageFolder+'/Rxn%s.jpg'%i,PlotFolder+'/Plot%s.jpg'%i, i)

led.off()

led2.off()

led3.off()

camera.stop_preview()

DataFile.close()

plt.plot(Time, Slot1,'co-', label=Sample1)

plt.plot(Time, Slot2,'ms--',label=Sample2)

plt.plot(Time, Slot3, 'g^:', label=Sample3)

plt.legend(loc=7, frameon=False)

plt.xlabel('Time')

plt.ylabel('Intensity (Au)')

plt.title('Fluorescence Monitoring in Vacuum Chamber')

plt.savefig('Fluorescence Monitoring in Vacuum Chamber')

plt.close()

Supplementary Code 2. Python code for reanalyzing collected images.

\# -*- coding: utf-8 -*-

"""

Created on Tue Sep 24 18:19:38 2019

@author: aufde025

"""

######To reanalyze old images#######

import os

import numpy as np

from PIL import Image

import matplotlib.pyplot as plt

ImageFolder=input('Name of folder where Rxn Images are:')

path, dirs, files = next(os.walk(r"C:\Users\aufde025\Google Drive\EA-Lab_Sept2019\Nanomotor\7-26-19_dessication_run4_images"))

file_count = len(files)

Data=input('Name file to save pixel intensities:')

#insert the name of the file where the data is being stored once analyzed

DataFile=open(Data,'a+')

#######My function for image analysis############

def PixelAvg(image, Slot1, Slot2, Slot3):

#loading the image in:

im=Image.open(image)

#imarray=np.array(im)

#to take a subset from image for each tube

#Slot1=imarray[50:240,1:427]

#Slot2=imarray[52:240,427:854]

#Slot3=imarray[50:240,855:1279]

#to go from an array back to an image:

#this is only necessary if you need to view your subset:

#Slot1Im=Image.fromarray(Slot1)

#Slot2Im=Image.fromarray(Slot2)

#Slot3Im=Image.fromarray(Slot3)

#converting Images to grayscale:

Slot1Gray=np.array(im.convert('L'),'f')[50:240,1:427]

Slot2Gray=np.array(im.convert('L'),'f')[52:240,427:854]

Slot3Gray=np.array(im.convert('L'),'f')[50:240,855:1279]

#finding the index of Max value of the image

#this will use that bright dot at the bottom of the tube to base my pixel subset off of

Slot1MaxInd=np.unravel_index(np.argmax(Slot1Gray), (190,426))

Slot2MaxInd=np.unravel_index(np.argmax(Slot2Gray), (188,427))

Slot3MaxInd=np.unravel_index(np.argmax(Slot3Gray), (190,424))

#after finding the max value indexes above, taking the subset to average over:

Slot1Pixel=Slot1Gray[(Slot1MaxInd[0]-30):(Slot1MaxInd[0]-10),(Slot1MaxInd[1]-10):(Slot1MaxInd[1]+10)]

Slot2Pixel=Slot2Gray[(Slot2MaxInd[0]-30):(Slot2MaxInd[0]-10),(Slot2MaxInd[1]-10):(Slot2MaxInd[1]+10)]

Slot3Pixel=Slot3Gray[(Slot3MaxInd[0]-30):(Slot3MaxInd[0]-10),(Slot3MaxInd[1]-10):(Slot3MaxInd[1]+10)]

#now find the mean of your pixel selection:

S1mean=np.mean(Slot1Pixel)

S2mean=np.mean(Slot2Pixel)

S3mean=np.mean(Slot3Pixel)

Slot1.append(S1mean)

Slot2.append(S2mean)

Slot3.append(S3mean)

return(S1mean, S2mean, S3mean, Slot1, Slot2, Slot3);

Slot1=[]

Slot2=[]

Slot3=[]

values=[]

for i in range(file_count):

values=PixelAvg(ImageFolder+'/Rxn%s.jpg'%i, Slot1, Slot2, Slot3)

DataFile.write(str(values[0])+'\t'+str(values[1])+'\t'+str(values[2])+'\n')

values=[]

DataFile.close()

Supplementary Code 3. Python code for generating fluorescence plot overlays for existing image sets.

\# -*- coding: utf-8 -*-

"""

Created on Tue Sep 24 10:14:15 2019

@author: aufde025

"""

import csv

import matplotlib.pyplot as plt

import os

from PIL import Image

#####Code to make the new graphs##########

Sample1=input('Name of sample in Slot 1:')

Sample2=input('Name of sample in Slot 2:')

Sample3=input('Name of sample in Slot 3:')

PlotFolder=input('Name a folder to save plot images:')

ImageFolder=input('Name a folder where rxn images are:')

OverlayFolder=input('Name a folder to save new overlay images:')

Title=input('What do you want your graph title:')

Interval=float(input('How often did your script take photos (in mins):'))

pause=input('Press enter when you are ready to begin')

os.makedirs(PlotFolder)

#####function to make overlays#######

def ImageView(TubePic,PlotPic,i):

background= Image.open(TubePic)

overlay= Image.open(PlotPic)

#added dpi to plt saving so I no longer need to resize

#wsize=int(min(background.size[0], background.size[1])*0.75)

#wpercent=(wsize/float(overlay.size[0]))

#hsize=int((float(overlay.size[1])*float(wpercent)))

#simage=overlay.resize((wsize, hsize))

#mbox=background.getbbox()

#sbox=simage.getbbox()

#box=(mbox[2]-sbox[2],mbox[3]-sbox[3])

background.paste(overlay, (400,360))

background.save(OverlayFolder+'/Ovly%s.jpg'%i, 'jpeg')

#background.show(background.thumbnail((700,600))) #this resizes it so it fits on Raspi screen

return(background);

#######code to make new graph#########

Time=[]

i=0

mydpi=100

val1=[]

val2=[]

val3=[]

with open('7-26-19_dessication_run4_intensities') as csvfile:

readCSV=csv.reader(csvfile,delimiter='\t')

for row in readCSV:

Time.append(i*Interval)

val1.append(float(row[0]))

val2.append(float(row[1]))

val3.append(float(row[2]))

plt.plot(Time, val1,'g-', label=Sample1)

plt.plot(Time, val2,'r--',label=Sample2)

plt.plot(Time, val3, 'k^:', label=Sample3)

plt.legend(loc=7, frameon=False)

plt.xlim(0,Time[i]+5)

plt.ylim(0,50)

plt.xlabel('Time')

plt.ylabel('Intensity (Au)')

plt.title(Title)

plt.savefig(PlotFolder+'/Plot%s.jpg'%i, dpi=mydpi*0.75)

i=i+1

ImageView(ImageFolder+'/Rxn%s.jpg'%i,PlotFolder+'/Plot%s.jpg'%i, i)

plt.close()

Supplementary Table 1. Thermal midpoints of **G4Redox.** Measurements were obtained by UV-vis monitoring of A­_260_ and A­_295_.

|  | T_M_ (°C) | |
| --- | --- | --- |
| [NaClO_4_] (M) | duplex | G4 |
| 0.1 | 70.5 | High |
| 1 | 73.6 | High |
| 2 | 66.0 | 75.4 |
| 3 | 56.9 | 69.7 |
| 4 | 47.9 | 66.7 |
| 5 | 35.5 | 60.8 |
| 6 | 29.8 | 58.8 |
| 7 | Low | 53.4 |
| 8 | Low | 52.5 |
| Saturated (ca. 9 M) | Low | 51.8 |

Supplementary Figure 37. Comparative kinetic measurements of oxidation of amplex red by **G4Redox** and **G4-SwitchR** in the presence of varying concentrations of NaClO_4_.

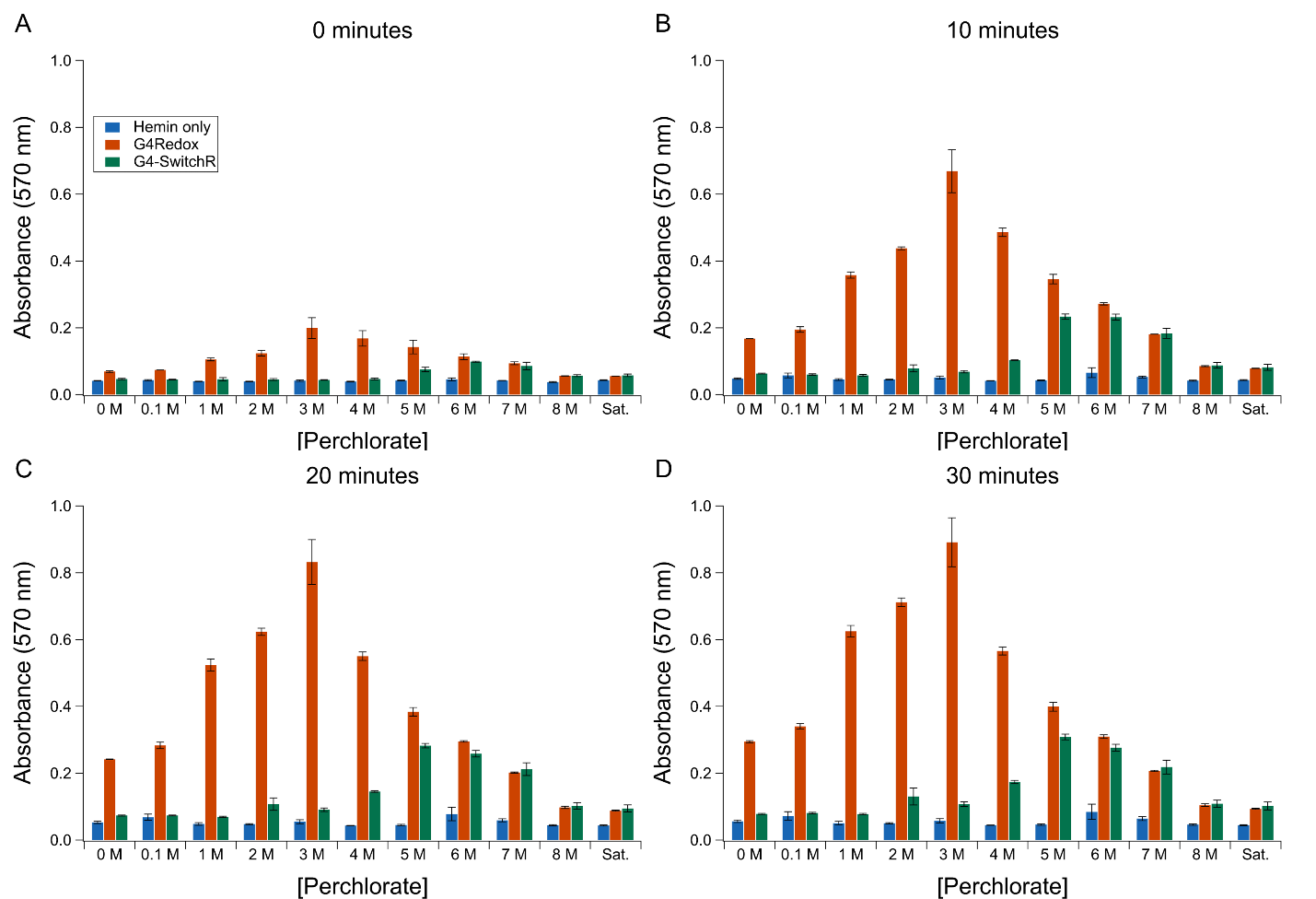

Supplementary Figure 38. A_295_-monitored melting curve of the **G4-SwitchR** system.

Melt samples consisted of 5 µM **G4Redox** and 5 µM **G4Redox-Comp** in 5 mM sodium phosphate buffer, pH 7.4. A_295_ increases with G4 formation. The trace initially starts at low values when **G4Redox** is duplexed with **G4Redox-Comp**, increases between ca. 20 and 40 °C as the duplex separates and **G4Redox** forms a G-quadruplex, then decreases between ca. 40 and 70 °C as the **G4Redox** G-quadruplex unfolds.

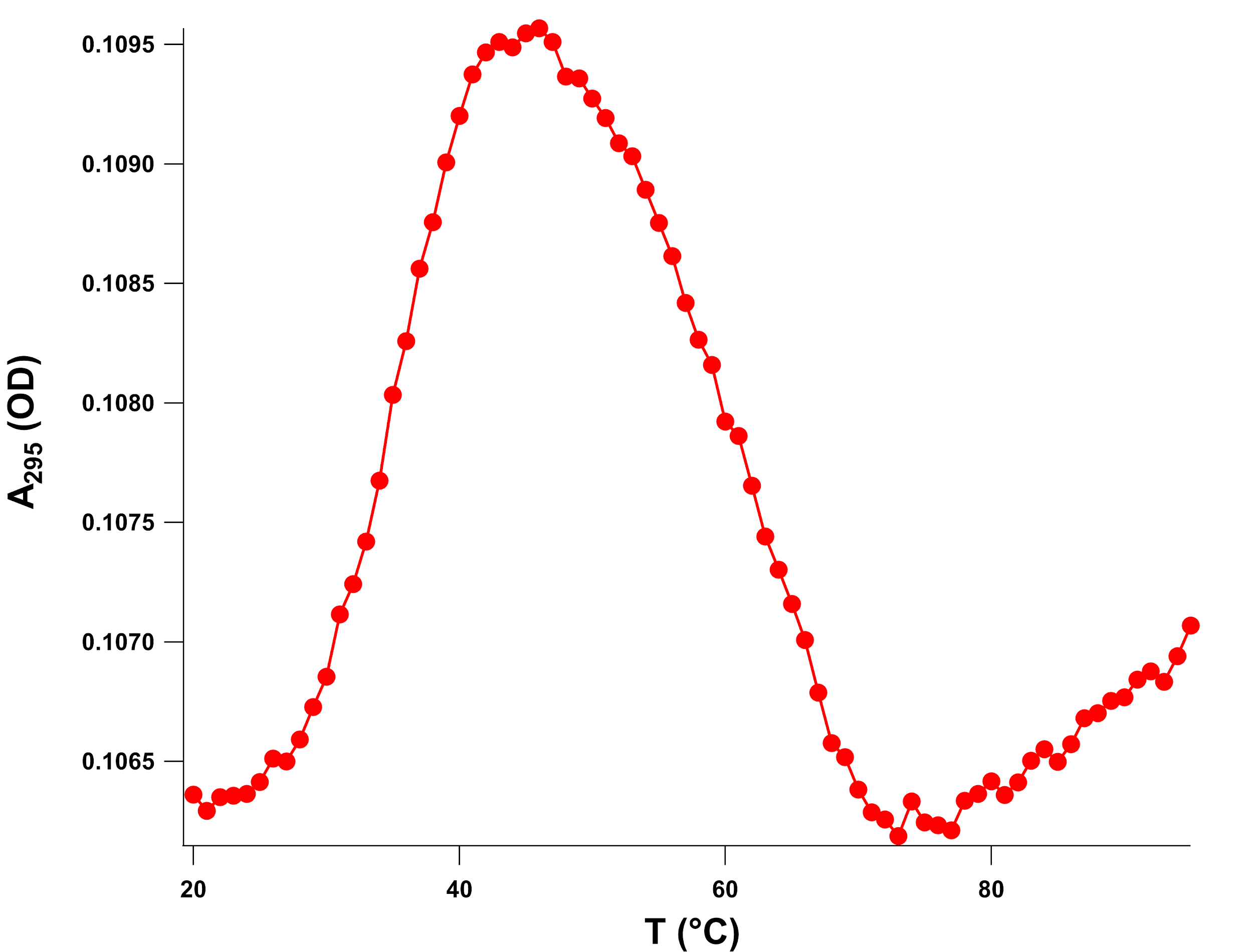

Supplementary Figure 39. A_260_-monitored melting curve of the **G4-SwitchR** system.

A_260_, which increases with unstacking of bases, exhibits the same transitions as the A_295_ trace. A_260_ initially starts at low values when **G4Redox** is duplexed with **G4Redox-Comp**, increases between ca. 20 and 40 °C as the duplex separates and **G4Redox** forms a G-quadruplex. A_260_ then increases further between ca. 40 and 70 °C as the **G4Redox** G-quadruplex unfolds. Melt samples consisted of 5 µM **G4Redox** and 5 µM **G4Redox-Comp** in 5 mM sodium phosphate buffer, pH 7.4.

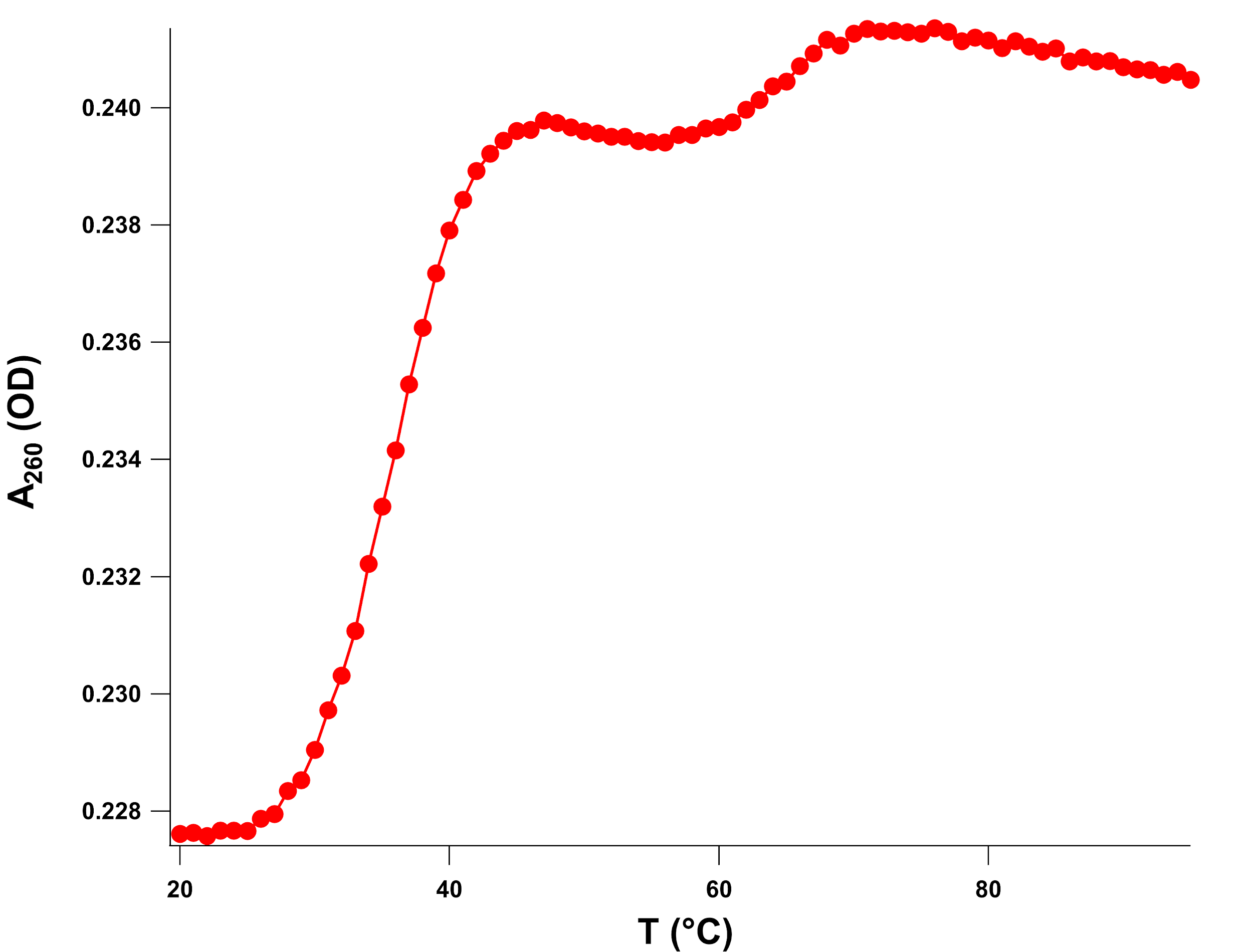
